# Activation of the endoplasmic reticulum stress regulator IRE1α compromises pulmonary host defenses

**DOI:** 10.1101/2024.09.28.615286

**Authors:** Amit Sharma, Linda M. Heffernan, Ky Hoang, Samithamby Jeyaseelan, William N. Beavers, Basel H. Abuaita

## Abstract

The endoplasmic reticulum (ER) stress sensor inositol-requiring enzyme 1-α (IRE1α) is associated with lung infections where innate immune cells are drivers for progression and resolution of inflammation. Yet, the role of IRE1α in pulmonary innate immune host defense during acute respiratory infection remains unexplored. Here, we found that activation of IRE1α in infected lungs compromises immunity against methicillin-resistant *Staphylococcus aureus* (MRSA)-induced primary and secondary pneumonia. Moreover, activation of IRE1α in MRSA-infected lungs and alveolar macrophages (AMs) leads to exacerbated production of inflammatory mediators followed by cell death. Ablation of myeloid IRE1α or global IRE1α inhibition confers protection against MRSA-induced pneumonia with improves survival, bacterial clearance, cytokine reduction, and lung injury. In addition, loss of myeloid IRE1α protects mice against MRSA-induced secondary to influenza pneumonia by promoting AM survival. Thus, activation of IRE1α is detrimental to pneumonia and therefore, it shows potential as a target to control excessive unresolved lung inflammation.

## Introduction

The endoplasmic reticulum (ER) stress response has been associated with many inflammatory lung diseases including respiratory infections.^1^ The ER serves as an essential organelle where biosynthesis and sorting of lipids and proteins, especially secreted molecules, takes place.^2^ Additionally, the ER is a major site for calcium storage, which can be released to control numerous biological processes. During infection, cells are exposed to environmental stressors, such as hypoxia and nutrient deprivation,^3,4^ and produce inflammatory mediators important for immune defense. However, these events often lead to overload and impair proper functioning of the ER.^5^ Halting any ER functions causes stress that leads to the activation of an adaptive response termed the unfolded protein response (UPR).^6,7^ The UPR is a signaling network that aims to restore cellular homeostasis by increasing protein folding and degradation of misfolded proteins, as well as reducing *de novo* protein synthesis and modulating calcium storage capacity. Many microbial pathogens, including respiratory viruses and bacteria, reportedly activate the UPR in the lung.^8–10^ A majority of previous studies focused on understanding the contribution of UPR signaling to viral pathogenesis. Viruses such as influenza and SARS-CoV-2 exploit UPR activity to enhance their pathogenesis, while pathogenesis of RSV (respiratory syncytial virus) is impaired by UPR activation through ill-defined mechanisms.^9–11^ However, it remains unclear whether the UPR aids or impedes immune defenses against the acute phase of bacterial respiratory infection.

In mammals, the UPR is controlled by three resident ER sensors, of which inositol-requiring enzyme 1 alpha (IRE1α) is most associated with downstream activation of immune cells. ^12,13^ IRE1α has a sensor domain that recognizes unfolded proteins in the ER lumen and enzymatic domains (kinase and endoribonuclease) that initiate UPR signaling in the cytosol.^7^ Once cells encounter stressors, the sensor domain of IRE1α initiates its oligomerization, autophosphorylation, and activation of cytoplasmic enzymatic domains. The endoribonuclease of IRE1α then splices *Xbp1* mRNA to allow translation of the transcription factor X-box Binding Protein-1 (Xbp1), ^14–16^ which induces expression of many genes that improve ER protein folding and degradation.^17,18^ The IRE1α endoribonuclease also regulates the expression of many UPR targets through post-transcriptional modifications via a process termed *regulated IRE1α-dependent decay* (RIDD) to restore ER homeostasis.^19,20^ Further, in addition to self-phosphorylation, the kinase domain of IRE1α can directly phosphorylate other host factors, such as TRAF-2, which activates MAPK signaling including activation of c-Jun N-terminal kinase (JNK).^21^ Our previous work and that of others suggests that infection of innate immune cells by bacterial pathogens engages the IRE1α pathway through Toll-like receptor (TLR)/MyD88 signaling.^13,22^ Importantly, activation of IRE1α by TLR ligation promotes expression of inflammatory genes including pro-inflammatory cytokines (TNF-α, IL-1β, and IL-6).^13,23–25^ Since inflammation is both necessary for immunity as well as pathogenic when occurring in excess or at inappropriate times, defining the role of innate immune IRE1α in respiratory infections is essential.

Methicillin-resistant *Staphylococcus aureus* (MRSA) infection is a leading hospital-borne infection that causes a wide spectrum of diseases ranging from localized skin infection to severe pneumonia and sepsis.^26^ *Staphylococcus aureus* is also a common causative agent of ventilator-associated pneumonia in intensive care units.^27^ Our previous studies have demonstrated that IRE1α is required for immunity against localized MRSA skin infection,^22,25^ where macrophages and neutrophils are essential effector immune cells. Further, we found that MRSA infection stimulates IRE1α activation in peritoneal and bone-marrow derived macrophages to increase production of pro-inflammatory cytokines.^22,24^ A recent study demonstrated that macrophage IRE1α is essential for production of prostaglandin E_2_ (PGE_2_) during inflammation to promote chronic pain.^28^ However, macrophages from different tissues mount variable magnitudes of inflammatory responses to promote host defense without disrupting tissue-specific functions.^29^ In the lung, PGE_2_ specifically acts as an anti-inflammatory molecule promoting tissue repair,^30^ suggesting that IRE1α may differentially control the tissue-specific function of lung macrophages.

Here, we investigated the direct contribution of IRE1α to the inflammatory responses of alveolar macrophages (AMs) *in vitro* during MRSA infection and of myeloid IRE1α to MRSA-induced primary and secondary pneumonia *in vivo*. Our results demonstrate that signaling by the IRE1α-Xbp1 pathway of the UPR imparts excessive AM inflammatory responses, which compromises immunity against MRSA-induced primary and secondary pneumonia. Moreover, the findings in this study suggest the potential for modulation of IRE1α signaling to promote the resolution of acute inflammation in the lungs.

## Results

### Activation of the IRE1α-Xbp1 pathway in AMs during MRSA infection promotes production of inflammatory mediators

Activation of IRE1α occurs in many lung inflammatory diseases where AM functions are critical for disease progression and resolution.^1^ To define the role of IRE1α in the AM inflammatory response, we first assessed IRE1α activation during bacterial infections. We chose methicillin-resistant *Staphylococcus aureus* (MRSA) as an infectious microbe since MRSA is a common causative agent of hospital-acquired pneumonia.^31^ AMs were infected with MRSA and IRE1α activation was assessed by quantifying the spliced form of *Xbp1* at 6 hours post-infection (hpi). MRSA-infected AMs increased the level of spliced *Xbp1*, which was completely abolished when AMs were treated with the IRE1α endoribonuclease inhibitor 4µ8C (Figures 1A and 1B). These data indicate that AM IRE1α is activated during MRSA infection.

**Figure 1.**
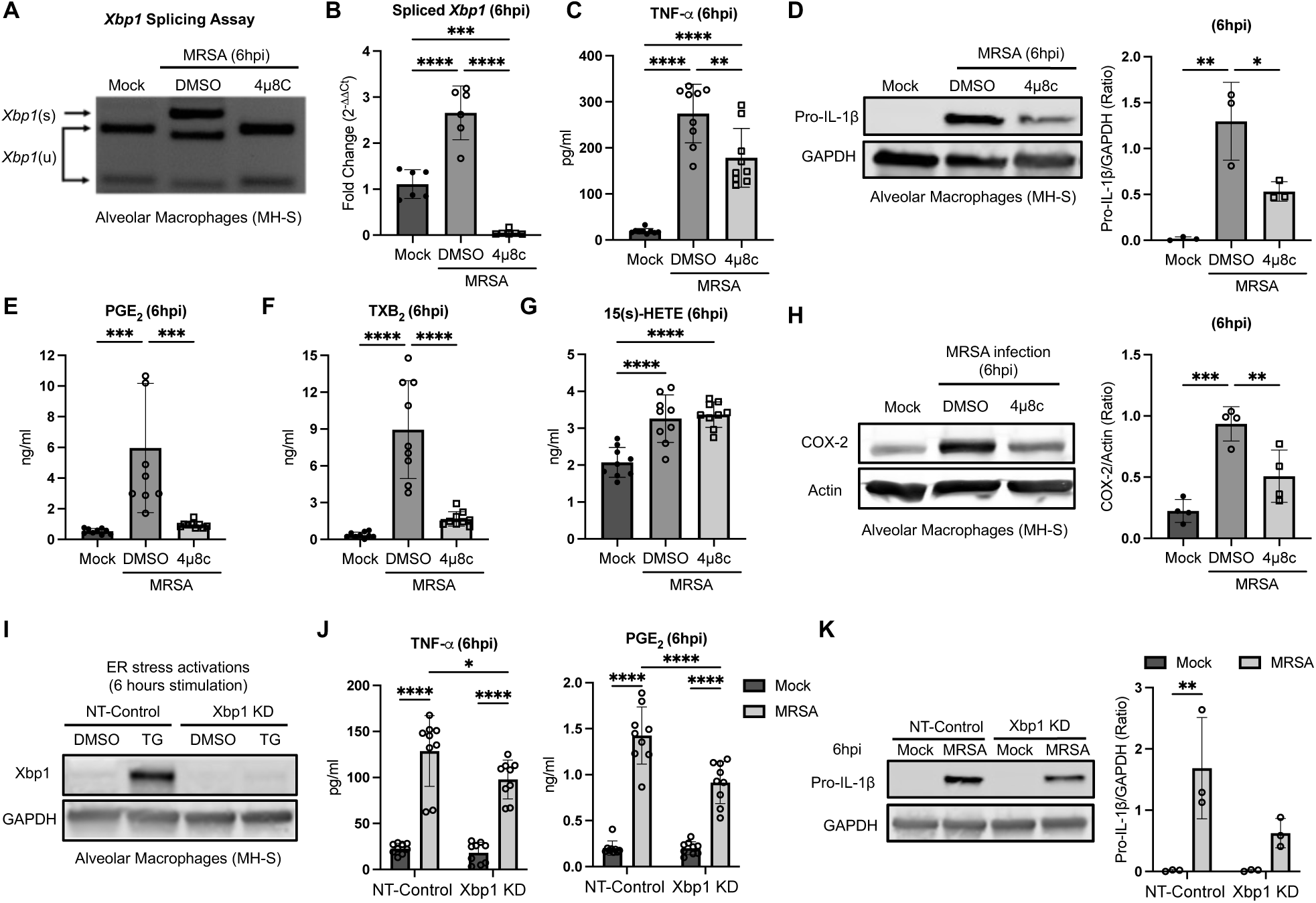
IRE1α-Xbp1 pathway is activated in AM and enhances production of pro-inflammatory mediators. (A) Representative image of DNA agarose gel of *Xbp1* splicing assay. Cells were left untreated (Mock) or infected with MRSA in the presence or absence of IRE1α endonuclease inhibitor (4µ8C). *Xbp1* RT-PCR product was digested with *PstI* restriction enzyme to cleave the unspliced transcript. The digested PCR products yield two smaller fragments representing unspliced (U) *Xbp1* and one larger fragment representing spliced (S) *Xbp1.* (B) The expression level of spliced *Xbp1* was quantified by RT-qPCR relative to *Gapdh*. (C) TNF-α level in culture media of AMs was quantified by ELISA at 6 hpi. (D) Immunoblot analysis of pro-IL-1β and GAPDH from cell lysates of AMs. Levels of protein expression were quantified by analyzing band intensity using Image J software. Graph indicates the means ± SD of pro-IL-1β expression levels relative to GAPDH. (E-G) Levels of eicosanoids; PGE_2_ (E), TXB_2_ (F), and 15(s)HETE (G) in culture media of AMs were quantified by ELISA at 6 hpi. (H) Immunoblots analysis of COX-2 protein expression of AMs. COX-2 expression was quantified relative to Actin by analyzing band intensity using Image J software. (I) Representative immunoblots of Xbp1 protein expression from NT-control and Xbp1 KD AMs when cells were treated with DMSO (control) or stimulated with an ER stressor Thapsigargin (TG, 1 µM) for 6 h. (J) Levels of TNF-α and PGE_2_ in culture media of NT-control and Xbp1 KD AMs. (K) Representative immunoblots of pro-IL-1β and GAPDH from cell lysates of NT-control and Xbp1 KD AMs when cells were untreated (Mock) or infected with MRSA for 6 h. Levels of protein expression were quantified as in panel D. Graphs indicate the mean of n≥3 independent experiments ± SD.

To define the functional consequences of IRE1α activation on AM inflammatory responses, we first monitored the production of inflammatory cytokines (TNF-α and pro-IL-1β) and lipid mediators (PGE_2_, TXB_2_, and 15(s)-HETE) when AMs were infected with MRSA in the presence or absence of IRE1α inhibitor (Figures 1C-1G). MRSA infection triggered a robust immune response by inducing the production of each of these inflammatory mediators, and inhibition of IRE1α reduced the production of TNF-α, pro-IL-1β, PGE_2_, and TXB_2_. IRE1α control of the production of inflammatory lipid mediators was not applicable for all eicosanoids as 15(s)-HETE was induced during infection but was not affected by inhibition of IRE1α (Figure 1G). In addition, the levels of lipoxygenase-dependent eicosanoids, LTB_4_ and LTC_4_, in culture supernatants were neither significantly increased during infection nor changed in the presence of 4µ8C (Figure S1A). Because PGE_2_ and TXB_2_ are produced via the cyclooxygenase (COX-2)-dependent eicosanoid pathway while 15(s)-HETE, LTB_4_, and LTC_4_ are all downstream of lipoxygenase,^32^ we monitored COX-2 expression in MRSA-infected AMs. Expression of COX-2 protein was increased during MRSA infection relative to uninfected cells, and IRE1α inhibition reduced COX-2 expression by infected AMs (Figure 1H). IRE1α-mediated activation of the cyclooxygenase pathway occurred by increasing transcription levels of *Ptgs2* (*Cox-2)* and *Ptges*, but not *Ptgs1* (*Cox-1)* (Figure S1B). By contrast, MRSA infection reduced expression of lipoxygenase (*Alox15*, *Alox5*, and *Alox5ap*) independent of IRE1α inhibition (Figure S1C), indicating that IRE1α controlled the production of PGE_2_ and TXB_2_ by reducing the expression of selected genes in the cyclooxygenase pathway. We next investigated whether IRE1α enhances cytokine productions via activation of the main downstream target, Xbp1. We generated a stable *Xbp1* knockdown AM (Xbp1 KD) and non-target control (NT-control) cell lines by expression of shRNA-specific for *Xbp1* or non-target control via RNA interference (RNAi) lentivirus-based technology. Stimulating these cells with the potent ER stressor Thapsigargin (TG) induced Xbp1 protein expression in NT-control AMs, but not in Xbp1 KD AMs, indicating that *Xbp1* expression is silenced (Figure 1I). Importantly, silencing *Xbp1* phenocopied IRE1α inhibition and resulted in impairment of cytokine (TNF-α, pro-IL-1β) production and COX-2-dependent inflammatory mediators (PGE_2_) when compared to MRSA-infected NT-control AMs (Figures 1J and 1K). Collectively, our data suggest that the IRE1α-Xbp1 pathway is activated in MRSA-infected AMs and promotes production of inflammatory mediators including COX-2-dependent eicosanoids.

### AMs and neutrophils are essential in immunity against MRSA-induced pneumonia

To investigate the role of IRE1α in pulmonary innate immune host defense, we first established a MRSA-induced pneumonia infection model by inoculating wild-type C57BL/6J mice with 10^8^ colony-forming units (CFU) of MRSA via the oropharyngeal route to introduce the bacteria into the lung.^33^ IRE1α activation and lung tissue damage were assessed at 48 hpi. Mice infected with MRSA exhibited increased *Xbp1* splicing, indicating that IRE1α is activated in MRSA-infected lungs (Figure 2A). MRSA infection induced a robust lung edema, which was quantified based on the accumulation of protein and DNA in the bronchoalveolar lavage fluid (BALF) (Figure 2B), and increased lung cell death, which was detected by terminal deoxynucleotidyl transferase dUTP nick end labeling (TUNEL) (Figure 2C). Induction of edema and lung cell death was accompanied by disruption of alveolar structure. This was apparent by the loss of type I epithelial cells in lung histology sections when labeled with anti-T1-α, a marker of alveolar surface/type I epithelial cells (Figure 2C),^34^ and by staining lung histology sections with hematoxylin and eosin (H&E) (Figure S2A). These results indicate that MRSA infection induces IRE1α activation and acute lung injury. We next sought to define the contribution of different innate immune cells (AMs, neutrophils, or monocytes) to pulmonary host defense against MRSA. Mice were inoculated with clodronate liposomes via the oropharyngeal route to deplete AMs or anti-Ly6G antibodies intravenously to deplete neutrophils. Mice that received clodronate liposomes had a lower number of AMs in their BALF and lungs (Figure 2D). AM depletion renders mice highly susceptible to non-lethal dosage (10^7^ CFU) of MRSA challenge as these mice exhibited a reduced survival rate and had higher bacterial counts in the BALF and lungs when compared to mice that received control liposomes (Figures 2E and 2F). Likewise, neutrophil depletion also increased susceptibility to MRSA-induced pneumonia. Mice that received anti-Ly6G antibody had lower neutrophil infiltration, also known as polymorphonuclear leukocytes (PMNs), in MRSA-infected lungs, reduced survival during MRSA challenges, and had higher bacterial counts in the lungs (Figures 2G-2I). Interestingly, neutrophil depletion provoked a compensatory increase in inflammatory monocyte (MONO) recruitment to the infected lungs (Figure 2G). However, this increase in the recruited inflammatory monocytes was insufficient to protect mice against MRSA challenges.

**Figure 2.**
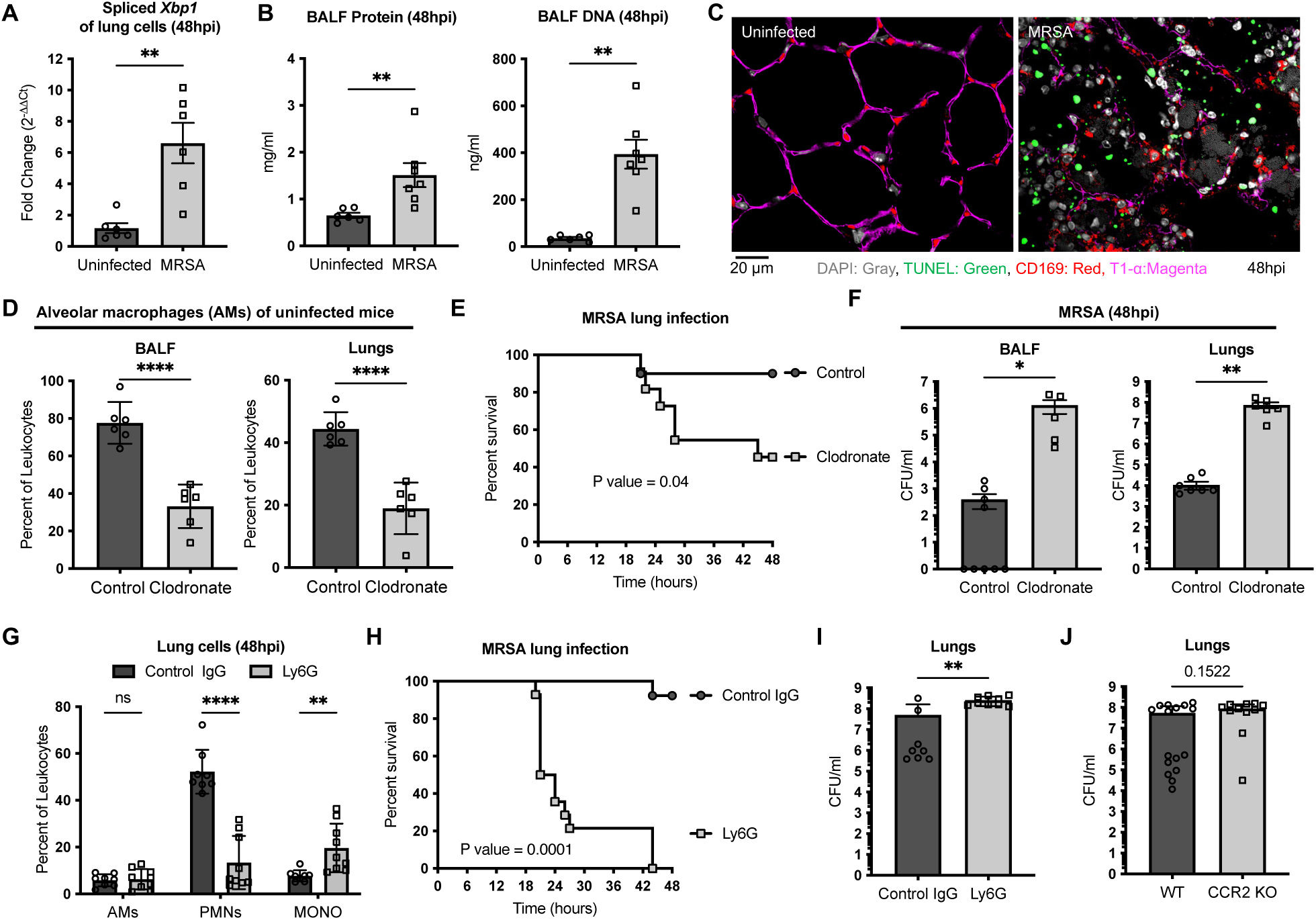
AMs and neutrophils are essential for immunity against MRSA-induced lung injury. (A) The levels of spliced *Xbp1* in lungs was assessed by RT-qPCR in uninfected mice or in mice infected with MRSA for 48 h. (B) The levels of protein and extracellular DNA in BALF of uninfected and MRSA-infected mice after 48 h. (C) Representative confocal microscopy images of lung histology sections from uninfected and MRSA-infected mice after 48 h. Histology sections were stained with TUNEL to label dead cells (green), anti-CD169 (red) to label macrophages, anti-T1α (magenta) to label type I epithelial cells, and DAPI (gray) to label DNA. (D) Flow cytometric analysis of AMs in BALF and lungs of mice receiving control liposomes or clodronate liposomes via oropharyngeal aspiration. Cells were stained with anti-CD45, anti-CD11c, and anti-CD170. (E) Survival of MRSA lung infected mice after aspiration of control liposomes or clodronate liposomes. (F) Bacterial burden in BALF and lungs of MRSA-infected mice after aspiration of control liposomes or clodronate liposomes. (G) Immunophenotyping of leukocytes in the lungs of mice when injected with control IgG or anti-Ly6G intravenously for two consecutive days prior to MRSA lung infection. Cells were stained for AM markers (anti-CD11c, anti-CD170), neutrophil (PMN) markers (anti-Ly6G, anti-CD11b), and monocyte (MONO) markers (anti-Ly6C, anti-CD11b) and analyzed by flow cytometry. (H, I) Survival (H) and bacterial burden (I) when mice were injected control IgG or anti-Ly6G intravenously for two consecutive days prior to MRSA lung infection. (J) Bacterial burden in lungs of wild-type (WT) and CCR2-knockout (CCR2 KO) mice infected with MRSA via oropharyngeal aspiration for 48 h. Survival rates were graphed using a Kaplan-Meier survival plot and p-values were calculated by Log-rank (Mantel-Cox) test. All other graphs represent the mean of n≥6 mice from at least two independent experiments ± SD.

Alternatively, to measure the requirement for inflammatory monocytes in pulmonary immunity against MRSA, we assessed the susceptibility of CCR2-deficient mice to MRSA-induced pneumonia. CCR2 signaling is required for monocyte recruitment during immune responses.^35^ CCR2-deficient mice exhibited a moderate increase in susceptibility to MRSA-induced pneumonia relative to wild-type mice only when mice were infected with a sub-lethal dose (10^8^ CFU) of MRSA. Moreover, CCR2-deficient mice exhibited reduced inflammatory monocyte recruitment to MRSA-infected lungs relative to wild-type mice (Figure S2B) and a subtle decrease in survival with an increase in lung MRSA burdens (Figures 2J and S2C). Together, our data establish that MRSA infection activates lung IRE1α and causes severe respiratory damage which may lead to death when there is ineffective cooperative pulmonary host innate immune defense by AMs, neutrophils, and monocytes.

### Ablation of myeloid IRE1α protects mice from MRSA-induced pneumonia

IRE1α is essential for placental development and thus, conventional IRE1α whole body knockout is lethal.^36^ Therefore, to define the role of IRE1α in pulmonary innate immune host response *in vivo*, we generated myeloid-specific IRE1α knockout mice by breeding *Ire1α^flox/flox^*mice with *Lyz2-Cre* mice that express Cre recombinase in myeloid cells.^37^ Analysis of IRE1α protein expression in isolated BALF AMs from mice showed IRE1α expression is restricted to AMs from control mice (IRE1α^f/f^), but not in the myeloid-IRE1α deficient mice (IRE1α^ΔMyeloid^) (Figure 3A). To address the role of IRE1α in pulmonary innate immune host defense, we infected IRE1α^Δmyeloid^ and control littermates (IRE1α^f/f^) with a sub-lethal dosage of MRSA via the oropharyngeal route and assessed survival, bacterial burden, leukocyte infiltration, and lung injury. After MRSA challenges, IRE1α^ΔMyeloid^ mice showed an enhanced survival rate over a 48 h period when compared to control IRE1α^f/f^ mice (Figure 3B), suggesting that deletion of myeloid IRE1α protects mice from MRSA-induced pneumonia. In line with the survival rate, we observed lower bacterial counts in BALF and lungs of IRE1α^ΔMyeloid^ mice relative to control IRE1α^f/f^ mice and reduced edema, which is indicated by the lower amounts of extracellular protein and DNA in BALF (Figures 3C and 3D). Immunophenotype analysis showed similar levels of neutrophils and monocytes recruited to BALF and lungs of MRSA-infected IRE1α^ΔMyeloid^ and IRE1α^f/f^ mice (Figure 3E). However, we observed higher AM counts in BALF of IRE1α^ΔMyeloid^ mice, which was also more evident when BALF cells were immobilized by cytospin and visualized using Diff-Quik staining (Figures 3E and 3F). These data suggest that deletion of myeloid IRE1α increases AM resilience to MRSA-induced cell death.

**Figure 3.**
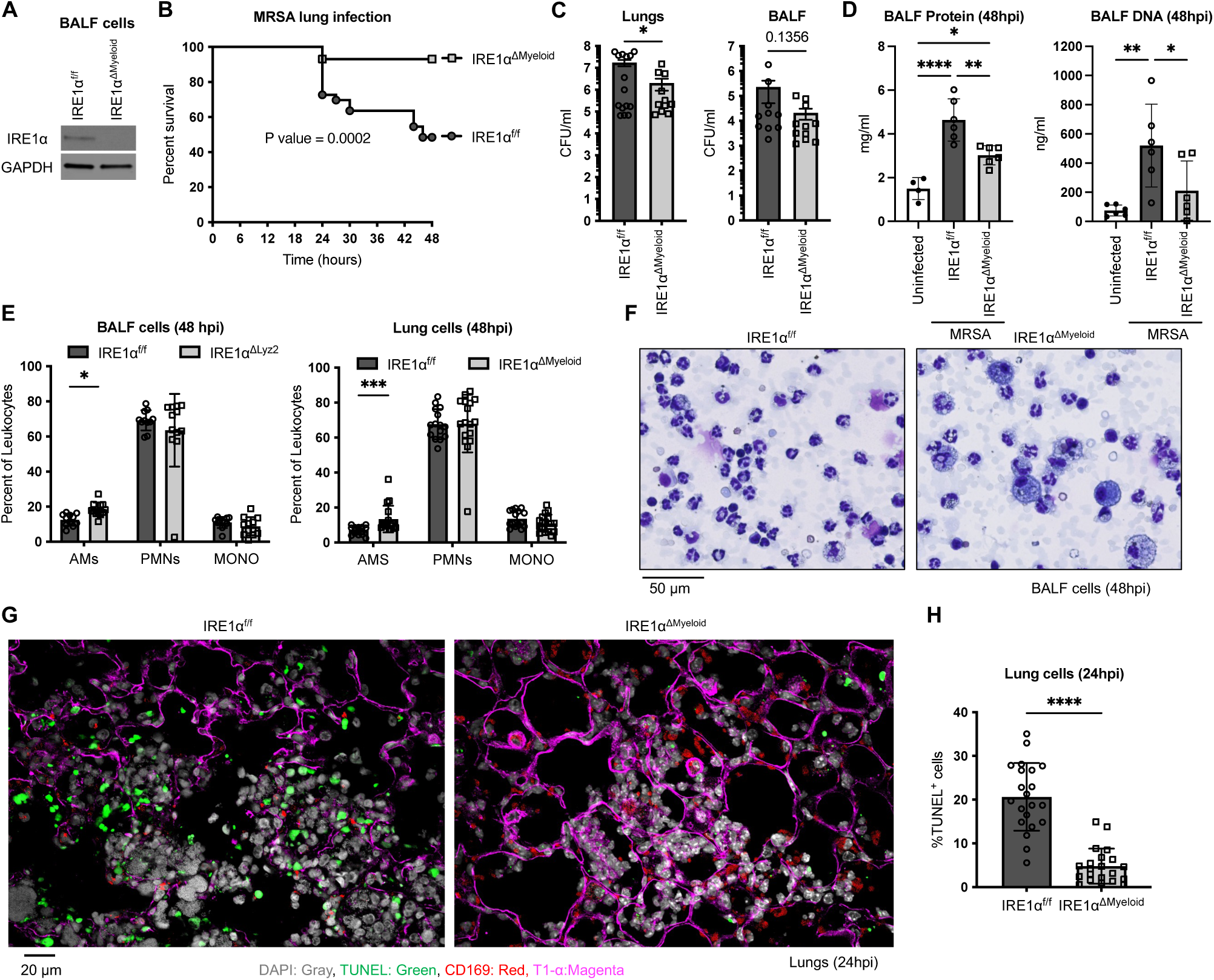
Silencing myeloid IRE1α protects mice from MRSA-induced pneumonia by inhibiting lung cell death. (A) Immunoblot analysis of IRE1α and GAPDH from AMs isolated from myeloid IRE1α-deficient (IRE1α^ΔMyeloid^) mice and littermate controls (IRE1α^f/f^). (B) Survival curves of IRE1α^ΔMyeloid^ and IRE1α^f/f^ mice infected with MRSA for 48 h. (C) Bacterial burden in BALF and lungs of MRSA-infected IRE1α^ΔMyeloid^ and IRE1α^f/f^ mice. (D) Protein and DNA levels in BALF of MRSA-infected IRE1α^ΔMyeloid^ and IRE1α^f/f^ mice. (E) Immunophenotyping analysis of leukocytes in BALF and lungs of IRE1α^ΔMyeloid^ and IRE1α^f/f^ mice infected via oropharyngeal aspiration for 48 h as in Figure 2G. (F) Representative images of BALF cells from MRSA-infected IRE1α^ΔMyeloid^ and IRE1α^f/f^ mice. (G) Representative confocal microscopy images of lung histology sections from IRE1α^ΔMyeloid^ and IRE1α^f/f^ mice when infected with MRSA for 24 hours. Sections were stained with TUNEL to label dead cells (green), anti-CD169 (red) to label macrophages, anti-T1α (magenta) to label type I epithelial cells, and DAPI (gray) to label DNA. (H) Percentage of TUNEL-positive cells in lung sections from IRE1α^ΔMyeloid^ and IRE1α^f/f^ mice. The percentage of TUNEL-positive cells was quantified relative to total cells based on DAPI staining by using Image J software. Graphs represent the mean of n≥12 IRE1α^ΔMyeloid^ mice and n≥11 IRE1α^f/f^ mice from three independent experiments.

To further characterize lung injury and cell death during MRSA-induced pneumonia, we evaluated lung histology sections of MRSA-infected mice at 24 hpi. Myeloid IRE1α deficient mice did not exhibit severe tissue injury based on H&E staining (Figure S3), and these mice only had mildly damaged alveoli in some of the lung lobes when assessed by TUNEL and anti-TI-α immunofluorescence staining (Figures 3G and 3H). In contrast, control wild-type mice exhibited severe lung injury, indicated by the presence of dense pockets of bacterial overgrowth and loss of type I epithelial cells lining some alveoli (Figure S3). In addition, TUNEL immunofluorescence staining showed more dead cells in the lungs of MRSA-infected control mice relative to myeloid IRE1α deficient mice (Fig. 3G and 3H). Together, our data suggested that myeloid IRE1α deficiency protects mice from MRSA-induced pneumonia by preventing excessive lung cell death.

### Innate immune IRE1α exacerbates pulmonary inflammation in response to MRSA-induced pneumonia

We next sought to characterize the contribution of myeloid IRE1α to the production of inflammatory mediators in MRSA-infected lungs. We used a bead-based multiplex assay to quantify the levels of cytokines and chemokines produced in the lungs during MRSA infection. We observed lower levels of several NF-κB-dependent inflammatory cytokines (TNF-α, IL-1β, and IL-6) and chemokines (MCP-1, KC, and MIP-1α) in MRSA-infected lungs of IRE1α^ΔMyeloid^ compared to lungs from control IRE1α^f/f^ mice (Figures 4A and 4B). It should be noted, a general reduction in all inflammatory mediators did not occur as several cytokines and chemokines were produced in similar amounts in the lungs of both mice (Figure S4A). Interestingly, we also observed a higher level of IFN-γ in the lungs of IRE1α^ΔMyeloid^ (Figure 4C), which may contribute to the observed protective response against MRSA infection.

**Figure 4.**
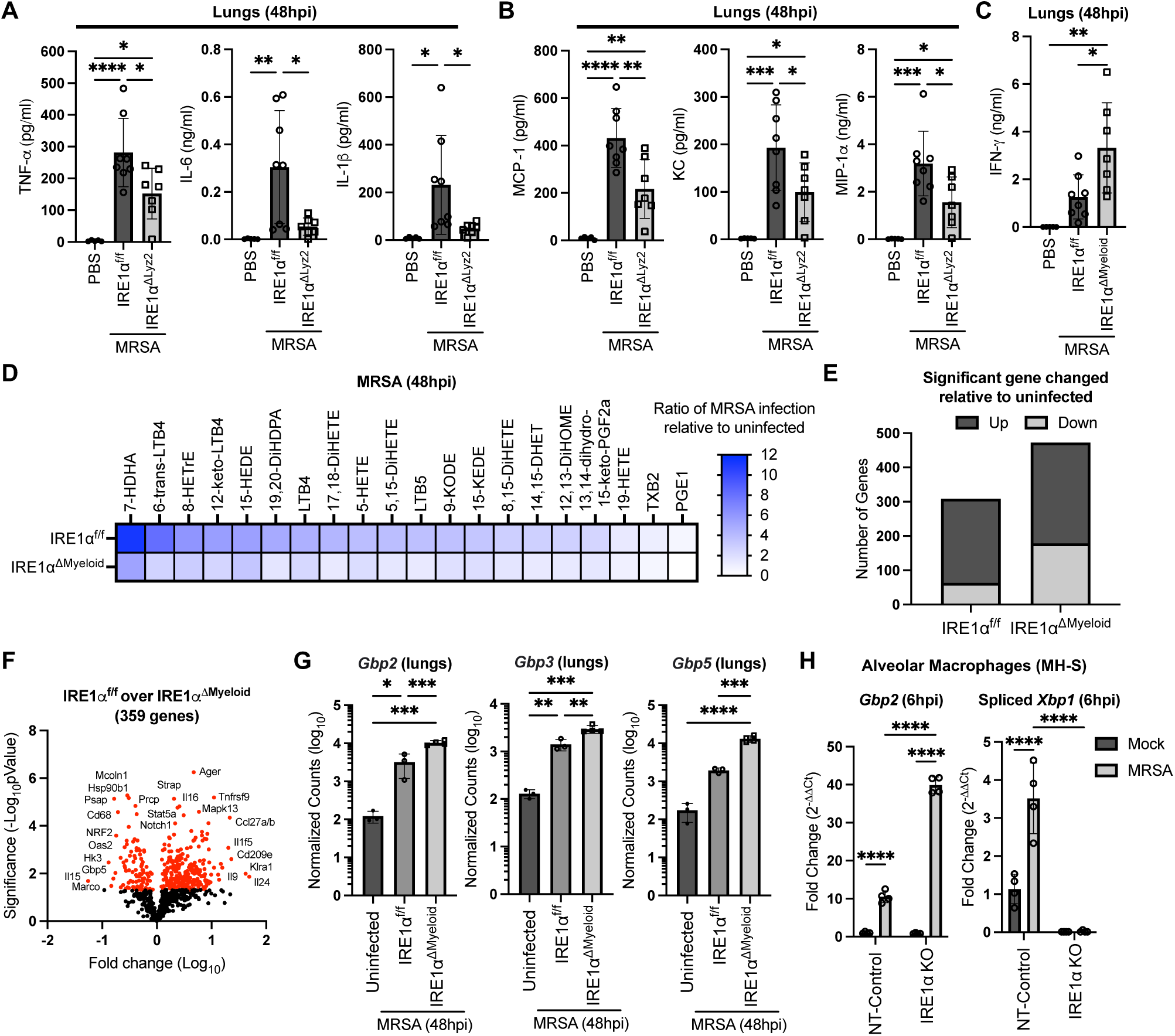
Myeloid IRE1α modulates pulmonary inflammatory responses during MRSA-induced pneumonia. (A-C) Levels of pro-inflammatory cytokines (A), chemokines (B), and IFN-γ (C) in supernatants of lung signal-cell suspensions from uninfected mice and MRSA-infected IRE1α^ΔMyeloid^ and control littermate IRE1α^f/f^ mice at 48 h. (D) Eicosanoid levels in supernatants of lung signal-cell suspensions from MRSA-infected IRE1α^ΔMyeloid^ and IRE1α^f/f^ mice were analyzed by mass spectrometry. Analytes are expressed as a ratio relative to uninfected mice. (E) Number of significantly changed genes in MRSA-infected lungs of IRE1α^ΔMyeloid^ and IRE1α^f/f^ mice relative to uninfected. Upregulated (dark gray) or downregulated (light gray) genes were identified based on log_10_ ratio of infected normalized counts to uninfected normalized counts. Significant genes were determined based on *P* value < 0.05. (F) Volcano plot of differentially expressed genes in MRSA-infected lungs from IRE1α^f/f^ mice over MRSA-infected lungs from IRE1α^ΔMyeloid^ mice. Red dots indicate genes that were significantly changed. (G) Quantification of interferon-inducible GTPase (*Gbp2, Gbp3,* and *Gbp5*) transcripts in lungs of uninfected mice, MRSA-infected IRE1α^f/f^, and MRSA-infected IRE1α^ΔMyeloid^ mice from NanoString data. (H) RT-qPCR of *Gbp2* and spliced *Xbp1* transcript levels of NT-control and IRE1α knockout (IRE1α KO) AMs when left untreated (Mock) or infected with MRSA for 6 h. Graphs represent the mean of n≥3 biological replicates ± SD.

To identify the contribution of myeloid IRE1α to the production of lipid immune mediators during MRSA challenge, we extracted lipids from the acellular fraction of dissociated lung single-cell suspensions and analyzed these by mass spectrometry. The levels of inflammatory lipids produced in MRSA-infected lungs were normalized relative to the levels in the lungs of uninfected mice (Figure 4D). Although prostaglandins were detected at lower levels in infected lungs at 48 h relative to uninfected controls, we observed reduced levels of PGE_1_, TXB_2_, and 15-keto-PGF_2a_ in lungs of MRSA-infected IRE1α^ΔMyeloid^ mice compared to control IRE1α^f/f^ mice. In addition, there were lower levels of other lipid mediators including leukotrienes (LTB_4_ and LTB_5_), HETEs (5-HETE, 19-HETE, and 8-HETE), and 7-HDHA produced in MRSA-infected lungs of IRE1α^ΔMyeloid^ compared to that in lungs of control IRE1α^f/f^ mice. The generally lower levels of lipid mediators produced in the lungs of MRSA-infected IRE1α^ΔMyeloid^ mice could be attributed to the lower levels of inflammation. These data suggest that myeloid IRE1α exacerbates lung inflammation during MRSA-induced pneumonia.

We next sought to characterize the contribution of myeloid IRE1α to the global pulmonary inflammatory response in MRSA-induced pneumonia. We used NanoString technology to quantify 785 transcripts of mouse inflammatory response genes in MRSA-infected lungs of IRE1α^ΔMyeloid^ and IRE1α^f/f^ mice as we performed previously.^38^ We found the number of inflammatory genes with expression that changed significantly in response to MRSA was higher in the lungs of IRE1α^ΔMyeloid^ mice than in the control IRE1α^f/f^ mice (Figure 4E). This was true for both the number of upregulated and downregulated genes. To define genes that are regulated by myeloid IRE1α during infection, we analyzed genes that differentially change in MRSA-infected IRE1α^f/f^ compared to MRSA-infected IRE1α^ΔMyeloid^ mice and identified 359 genes (Figure 4F and Table S1). This gene set contained IL-20 subfamily cytokines *Il19* and *Il24*, which were both increased in infected IRE1α^f/f^ relative to infected IRE1α^ΔMyeloid^ mice, indicating that myeloid IRE1α signaling is required for their induction in response to infection. In contrast, the antioxidant master regulator nuclear factor erythroid 2-related factor 2 (NRF2), immunometabolism regulator Hexokinase 3 (HK3), cluster of differentiation 68 (CD68), and macrophage receptor with collagenous structure (Marco) were among the genes increased in the absence of myeloid IRE1α. These data suggest that myeloid IRE1α signaling negatively regulates pulmonary oxidative stress via decreasing NRF-2 expression, immunometabolism via decreasing HK3 expression, and macrophage plasticity via decreasing M2 macrophage markers (CD68 and Marco). We then sorted the top 30 upregulated genes in infected IRE1α^f/f^ mice compared to infected IRE1α^ΔMyeloid^ mice. This set represents genes regulated by myeloid IRE1α during MRSA infection. This analysis revealed several inflammatory cytokines and chemokines (*Csf2*, *Csf3*, *Ccl20, Cxcl2*, *Cxcl5*, *Il17a*, *Il12a*, and others), which were increased in IRE1α^f/f^ and to a lesser extent in IRE1α^ΔMyeloid^ mice (Figure S4B). In this category, we also observed neutrophil-specific genes, such as *Lcn2* and *Elane*, with differentially increased expression in MRSA-infected IRE1α^f/f^ relative to IRE1α^ΔMyeloid^ mice. These results indicate that activation of myeloid IRE1α positively promotes MRSA-induced lung hyperinflammation. We next analyzed genes with increased expression in MRSA-infected IRE1α^ΔMyeloid^ relative to IRE1α^f/f^, which represent genes negatively regulated by IRE1α. This analysis identified several of IFN-γ-dependent inflammatory genes, including *Gbp2, Gbp5, Ifi203, Isg15, Oas1a, Oas2, Irf1, Irf7, Fcgr1, Fcgr4, and Stat-1*, which concurred with the increased IFN-γ production in MRSA-infected IRE1α^ΔMyeloid^ mice (Figure 4C). Importantly, several genes with expression increased in MRSA-infected IRE1α^ΔMyeloid^ relative to IRE1α^f/f^ are reported to have antimicrobial properties. Examples of such genes include *Gbp2, Gbp5, Ctsz, Ctss,* and *Psap* (Figures 4G, S4C, and Table S1), which may contribute to protection from MRSA-induced pneumonia observed in IRE1α^ΔMyeloid^ mice. To directly assess the contribution of loss of IRE1α signaling to the ability of alveolar macrophages to induce the antimicrobial interferon-induced guanylate-binding proteins, we quantified the expression of *Gbp2* in the CRISPR-Cas9 generated IRE1α knockout (KO) AMs relative to NT-control (Figure 4H). MRSA-infected IRE1α KO AMs failed to splice *Xbp1* transcripts, which validates deletion of IRE1α. Importantly, IRE1α KO AMs induced higher expression of *Gbp2* in response to MRSA when compared to NT-control AMs. Thus, the lack of IRE1α signaling increases expression of AM antimicrobial genes like *Gbp2* to promote MRSA clearance.

Lastly, we analyzed genes differentially downregulated during MRSA infection in IRE1α^f/f^ relative to IRE1α^ΔMyeloid^, aiming to identify IRE1α-dependent gene suppression during infection (Figure S4C). Genes downregulated in MRSA-infected IRE1α^f/f^, but not in IRE1α^Δmyeloid^, include *Ace, Cxcl15, Map3k1, Lrrk2, and Prcp*. This analysis also highlighted *Mcoln1, Hsp90b, NRF2, Ctsa, Gusb, and Ikbkg*, which were suppressed in MRSA-infected IRE1α^f/f^ and upregulated in MRSA-infected IRE1α^ΔMyeloid^. The contributions of these genes to protecting mice against MRSA-induced pneumonia will be investigated in our future studies. Collectively, our data suggest that ablation of myeloid IRE1α alters pulmonary inflammation into a protective response against MRSA.

### Global IRE1α inhibition dampens MRSA-induced pneumonia

To leverage our findings into a potential therapeutic approach, we sought to define whether inhibition of IRE1α endoribonuclease activity with the small molecule inhibitor, 4µ8C, would protect mice from MRSA-induced pneumonia. Mice were treated with 4μ8C or vehicle control via intraperitoneal injections 24 h prior to MRSA lung infection and then every 24 h throughout the duration of the experiment. IRE1α inhibition resulted in increased survival, similar to what was seen with myeloid IRE1α deficient mice, which was accompanied by reduced bacterial counts in BALF and lungs (Figures 5A and 5B). Mice treated with 4μ8C had similar levels of protein and DNA in BALF and had similar recruitment of neutrophils and monocytes to the BALF and lungs when compared to vehicle control (Figures 5C and 5D), thus, IRE1α inhibition did not alter vascular permeability in response to MRSA infection. Instead, the increased immune protection was associated with an increase in the relative abundance of alveolar macrophages (Figure 5D). To further visualize BALF cells from MRSA-infected mice treated with the IRE1α inhibitor, BALF cells were extracted, immobilized onto microscopy slides, and Diff-Quik stained (Figure 5E). Like MRSA-infected IRE1α^ΔMyeloid^ mice, mice treated with IRE1α inhibitor showed an increase in the relative abundance of macrophages in BALF during MRSA infection compared to BALF of infected control mice. Thus, this may suggest that the inhibition of IRE1α increases alveolar macrophage resilience in response to MRSA.

**Figure 5.**
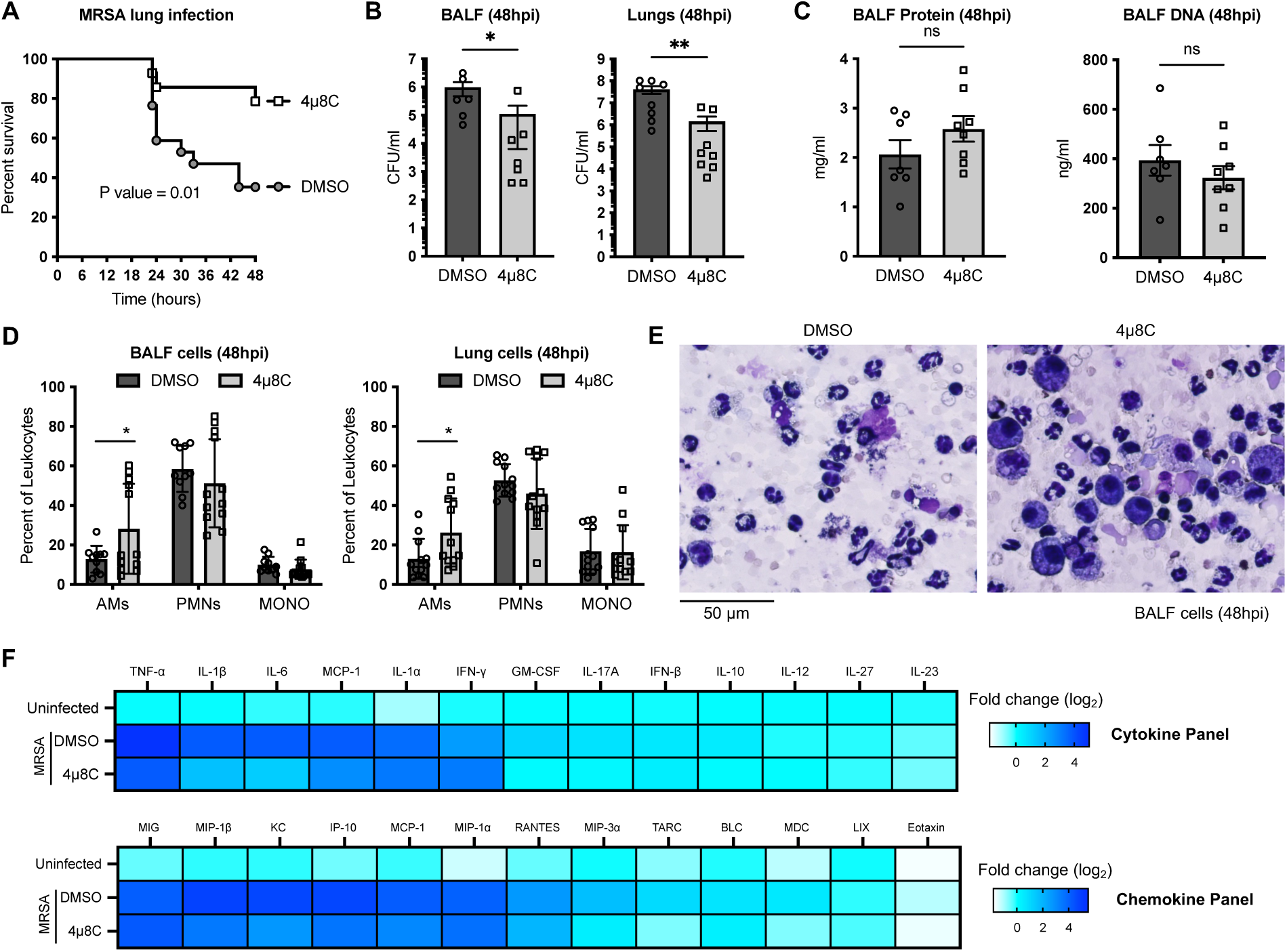
IRE1α inhibition confers protection against MRSA-induced pneumonia. (A) Survival of MRSA-infected mice when mice were treated with IRE1α inhibitor, 4µ8C, or control solvent. P value was calculated by the Log-rank (Mantel-Cox) test. (B-D) Bacterial burden (B), protein and DNA levels (C), and immunophenotyping analysis of leukocytes (D) in BALF and lungs of MRSA-infected mice when injected with 4µ8C or vehicle control via the intraperitoneal route. (E) Representative cytospin image of BALF cells from MRSA-infected mice treated with 4µ8C or vehicle control. (F) Heatmaps depicting expression of cytokines and chemokines in lungs of uninfected mice, MRSA-infected mice treated with 4µ8C, and MRSA-infected mice treated with vehicle control.

We next sought to determine whether IRE1α inhibition impacts production of inflammatory mediators in infected lungs. The lungs from infected mice treated with 4μ8C or vehicle control were dissociated into cell suspensions in phosphate buffered saline (PBS) and acellular supernatants were analyzed using bead-based flow cytometric immunoassay panels for cytokines and chemokines (Figure 5F). Mice treated with 4μ8C had lower amounts of cytokines (TNF-α, IL-1β, IL-1α) and chemokines (MIP-1β, KC, IP10, and MCP-1) than did controls. In contrast, IRE1α inhibition increased IFN-γ levels compared to vehicle control-treated mice, which phenocopies the ablation of myeloid IRE1α. Together, our data suggest that pharmacologically blocking IRE1α signaling protects mice from MRSA-induced pneumonia by reprogramming pulmonary inflammatory responses.

### Ablation of myeloid IRE1α protects mice against secondary MRSA pneumonia

Viral infections such as influenza and coronavirus substantially worsen secondary bacteria-induced pneumonia ^39,40^. A recent study showed that the unfolded protein response governed by IRE1α is among the most induced pathways in mice coinfected with influenza and MRSA.^41^ However, it is not clear whether ablation of myeloid IRE1α is protective against influenza/MRSA coinfection, as it is to primary MRSA infection. To establish influenza/MRSA coinfection, we first aimed to identify a viral dose and then a time point to introduce MRSA secondary to influenza. Mice were inoculated with 10 or 100 plaque-forming units (PFUs) of murine-adapted influenza A (PR8) via the oropharyngeal route. Survival and body weight were monitored over 12 days (Figures 6A and 6B). All mice started to lose weight by day 5 pi; however, only the mice infected with 100 PFU of PR8 exhibited mortality on day 8, while mice infected with 10 PFU eventually recovered. The data provides us with study conditions including an inoculum size of 100 PFU and a time point of day 6 post-influenza infection to induce lung damage sufficient for introduction of secondary MRSA infection without lethality. Interestingly, this time point is consistent with human clinical data, as secondary bacteria-induced pneumonia is typically observed within the first six days of influenza infection.^42^ When mice were challenged with a non-lethal dose of MRSA on day 6 post-influenza infection, all mice succumbed within 24 h while all mice infected with MRSA alone survived (Figures 6C and 6D). These data highlight that mice infected with influenza become highly susceptible to a non-lethal dosage of MRSA and establish a model for MRSA-induced secondary pneumonia.

**Figure 6.**
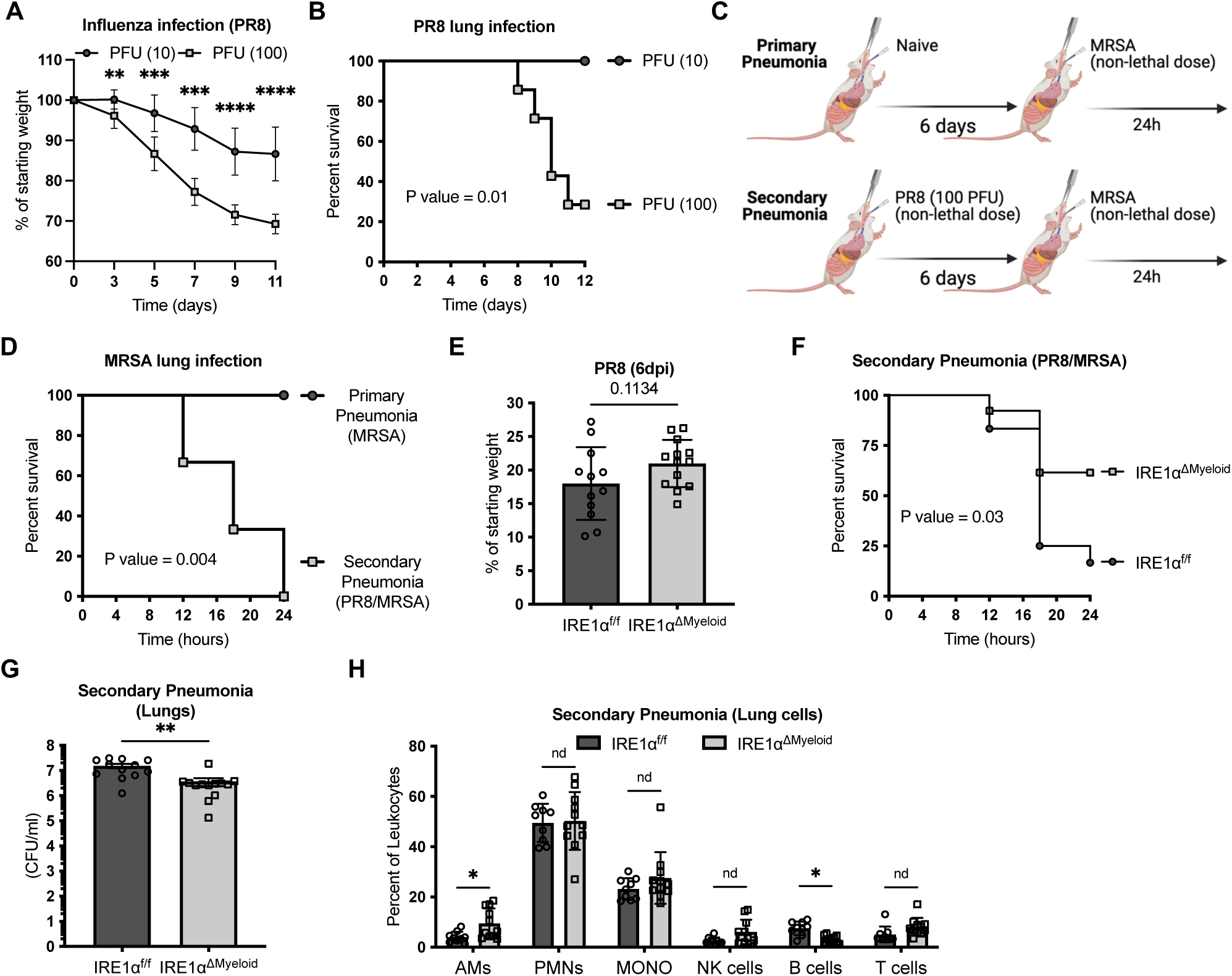
Myeloid IRE1α exacerbates influenza/MRSA coinfection. (A-B) Change in body weight (A) and survival (B) of mice infected with 10 or 100 PFU of PR8. Percentages were calculated relative to the starting bodyweight. (C) Schematic representation of MRSA-induced secondary pneumonia. (D) Survival curves of MRSA-induced primary and secondary pneumonia. (E) Body weight changes of IRE1α^ΔMyeloid^ and IRE1α^f/f^ mice infected with 100 PFU of PR8. (F-H) Survival (F), bacterial burden in lungs (G), and immunophenotyping of leukocytes in lungs (H) of MRSA-induced secondary pneumonia in IRE1α^ΔMyeloid^ and IRE1α^f/f^ mice at 24 h. Graphs in panels A, E, G, and H indicate the mean of n≥10 mice of each group from three independent experiments ± SD.

To investigate the role of myeloid IRE1α in MRSA-induced secondary pneumonia, IRE1α^ΔMyeloid^ mice and control littermates (IRE1α^f/f^) were infected with 100 PFU of PR8 for 6 days and then challenged with a non-lethal dose of MRSA (Figure 6C). Both IRE1α^ΔMyeloid^ and IRE1α^f/f^ mice lost a similar percentage of body weight in the first 6 days post-influenza infection, suggesting that a similar extent of lung injury occurred (Figure 6E). However, IRE1α^ΔMyeloid^ mice exhibited a higher survival rate post-MRSA challenge when compared to IRE1α^f/f^ mice (Figure 6F). The higher survival rate of IRE1α^ΔMyeloid^ mice coincided with greater MRSA clearance, as there were lower bacterial counts in the lungs of IRE1α^ΔMyeloid^ mice relative to lungs of IRE1α^f/f^ mice (Figure 6G). To further characterize the role of myeloid IRE1α in controlling the clinical outcomes of MRSA-induced secondary pneumonia, we analyzed the relative abundance of immune cells and production of inflammatory cytokines and chemokines. Similar to primary MRSA infection, there were higher AMs in influenza/MRSA coinfected IRE1α^ΔMyeloid^ mice relative to that in coinfected IRE1α^f/f^ mice (Figure 6H). Thus, this suggests that the lack of myeloid IRE1α also preserves AMs during coinfection. Remarkably, we also observed an altered relative abundance of lymphocytes in infected myeloid IRE1α deficient mice. There were fewer B cells and trending more T and NK cells in the lungs of influenza/MRSA coinfected-

IRE1α^ΔMyeloid^ mice relative to lungs of coinfected-IRE1α^f/f^ mice. The mechanism by which myeloid IRE1α regulates adaptive immune cell recruitment and function will be investigated in future studies. Lastly, we observed a similar level of inflammatory mediators produced in the lungs of IRE1α^ΔMyeloid^ and IRE1α^f/f^ mice during coinfection (Figure S5). This could indicate that the loss of myeloid IRE1α is insufficient to modulate lung inflammation during influenza/MRSA coinfection. Collectively, our data highlight that blocking the activity of myeloid IRE1α could be beneficial in protecting against bacteria-induced secondary pneumonia.

## Discussion

Acute respiratory infections are often associated with exuberant inflammation that can result in permanent lung damage and death. Activation of the ER stress sensor IRE1α can induce inflammation and its activation is associated with many lung diseases.^1^ However, whether IRE1α counteracts or supports the pathogenesis of bacteria-induced pneumonia is unknown. Because innate immune cells like alveolar macrophages and neutrophils are drivers for both the progression and resolution of respiratory infections, we aimed to characterize the role of innate immune IRE1α in MRSA-induced primary and secondary to influenza pneumonia. We found that ablation of myeloid IRE1α or global inhibition IRE1α activity protects mice from MRSA-induced primary and secondary pneumonia. MRSA lung infection causes pulmonary IRE1α activation, hyperinflammation, cell death, and lung injury resulting in mortality. Loss of IRE1α signaling tuned inflammatory responses toward type II interferons and away from NF-kB-mediated responses. Consequently, mice lacking myeloid IRE1 signaling exhibited less cell death and survived MRSA-induced pneumonia and MRSA-induced secondary to influenza pneumonia. These data highlight the potential benefit of modulating IRE1α activity as a novel therapeutic intervention to dampen lung inflammation and injury during acute respiratory infections.

Our findings support a model in which activation of innate immune IRE1α induces lung cell death that contributes to severity of pneumonia. The sub-lethal dose of MRSA used in our MRSA-induced pneumonia model caused death in approximately 50% of WT mice, while myeloid IRE1α-deficient mice were protected. MRSA-infected lungs appeared swollen and had several patches of cells possibly representing localized abscesses. Upon examination of lung histology sections, an enormous amount of cell death associated with loss of the organized airways was observed in WT mice and, to a significantly lesser extent, in myeloid IRE1α-deficient mice. Notably, we did not observe any bacteria or inflammatory cytokines in extrapulmonary organs like spleen and liver (data not shown). These data indicate that morbidity and mortality occur in mice due to the damaged alveolar sacs caused by the induction of massive lung cell death. A number of studies have shown that different mechanisms of programmed cell death, including inflammasome-mediated pyroptosis and receptor-interacting serine/threonine-protein kinase 1 (RIPK1)-dependent necroptosis, contribute significantly to the pathology of *S. aureus*-induced pneumonia.^43–46^ In addition, activation of inflammasomes increases inflammation and mortality in mice during influenza/*S. aureus* coinfection.^47^ While IRE1α can promote survival and cell fate, depending on the tissue and cell type, magnitude of stress and type of the stress, if ER stress is unresolved IRE1α acts as an executioner.^48^

Our work and that of others has shown that IRE1α activation induces and preserves thioredoxin-interacting protein (TXNIP), which in turn activates inflammasome-dependent pyroptosis.^24,49^ In addition, when ER stress is sustained, RIP1K can directly interact with IRE1α to induce necroptosis.^50^ However, it remains unclear how limiting myeloid IRE1α and cell death is beneficial. One possibility is that repressing IRE1α signaling preserves AMs in inflamed lungs to clear some of the dead cells and prevent high neutrophil activity in the lungs. Consequently, this would allow better oxygen exchange and prevent severe pneumonia and death. However, future experiments will be needed to determine the precise mechanism through which inhibition of IRE1a mediates these effects. Taken together, our data strongly suggest that IRE1α potentiates MRSA-induced pneumonia by enhancing pathways that lead to cell death perhaps via pyroptosis and/or necroptosis.

Silencing myeloid IRE1α signaling permits activation of type II interferon-mediated host defense to promote immunity against respiratory pathogens. Blocking IRE1α-Xbp1 signaling genetically or using a pharmacological approach attenuates production of several NF-κB-dependent pro-inflammatory mediators during MRSA infection both *in vitro* using murine alveolar macrophages and *in vivo* in infected lungs. Examples of these inflammatory mediators are TNF-α, IL-1β, IL-6, and PGE_2_. In contrast, we found that IFN-γ and IFN-γ-dependent antimicrobial genes, including *Gbp2, Gbp3 and Gbp5,* were among the upregulated genes when IRE1α signaling is inhibited. This indicates that myeloid IRE1α signaling can negatively control induction of the type II interferon response. Stimulation of immune cells with IFN-γ increased MRSA killing *in vitro*, which suggests that increased levels of IFN-γ in the lungs would enhance the ability of immune cells to kill respiratory pathogens like MRSA.^51^ However, in contrast to this, a recent study suggests that increasing levels of IFN-γ in response to influenza infection contributes to the extreme pathology of MRSA-induced secondary pneumonia by producing excessive TNF-α.^52^ In addition, mice lacking IFN-γ signaling (*Ifngr1* deficient mice) exhibit a similar susceptibility to primary MRSA-induced pneumonia as seen in WT mice.^53^ A possible explanation to this discrepancy is that primary MRSA infection does not produce IFN-γ in WT and *Ifngr1* deficient mice to drive IFN-γ-dependent host defense in the lungs. Both WT and *Ifngr1* deficient mice should trigger myeloid IRE1α signaling to repress production of IFN-γ in MRSA-infected lungs. Thus, presence and absence of *Ifngr1* is negligible to primary MRSA lung infection since the ligand (IFN-γ) is not made. In the context of MRSA-induced secondary pneumonia, the lack of myeloid IRE1α signaling could still reduce the negative impact of excessively produced TNF-α despite the high levels of IFN-γ. Another observation that remains unsettled is the source of IFN-γ produced in MRSA-infected lungs of myeloid IRE1α deficient mice. IRE1α could suppress IFN-γ production in myeloid-derived cells like AMs autonomously or indirectly through crosstalk between myeloid-derived cells and other immune cells such as NK and T cells. We and others have previously reported that activated macrophages can produce IFN-γ by an unknown mechanism.^38,54,55^ This implies that suppression of IRE1α signaling could be a mechanism by which AMs contribute to IFN-γ production in infected lungs. Alternatively, in the absence of IRE1α signaling, AMs could signal to other immune cells to stimulate and release IFN-γ. For example, our data suggest that myeloid IRE1α deficient mice exhibit a slight modulation in the abundance of B, T and NK cells in response to influenza/MRSA coinfection (Figure 6H), although it is unclear how this occurs. In our future studies, we will explore mechanisms by which myeloid IRE1α limits IFN-γ production and the practical significance to progression and resolution of different inflammatory diseases.

Tuning down production of lipid mediator PGE_2_ via inactivating myeloid IRE1α signaling promotes survival and may reduce chest pain associated with pneumonia. Production of PGE_2_ is elucidated during inflammation via the activity of COX-2.^56^ We observed that the IRE1α-Xbp1 axis selectively controls prostanoids (PGE_2_ and TXB_2_) through upregulation of COX-2 in murine alveolar macrophages and lungs upon MRSA infection. PGE_2_ can exhibit pro– and anti-inflammatory roles depending on which receptors it engages on target cells.^56^ In some contexts, PGE_2_ can suppress macrophage host defense by reducing phagocytosis and/or production of antimicrobial molecules like nitric oxide (NO).^57,58^ In addition, inhibition of COX-2 had a beneficial suppressive effect in hyperinflammatory lung diseases,^59^ inferring that modulation of PGE_2_ by inhibition of IRE1α signaling could mitigate disease severity. Interestingly, a recent study suggests that activation of IRE1α-Xbp1 promotes pain sensation via increasing production of PGE_2_.^28^ One of the main complications in patients with bacteria-induced pneumonia is severe chest pain, which could be driven in part by IRE1α-dependent production of PGE_2_. Therefore, we speculate that inhibition of IRE1α could have a dual advantage by reducing lung inflammation and relieving chest discomfort associated with pneumonia.

Recurrent activation of IRE1α following repeated exposure to similar or different stressors could be deleterious. Lungs are constantly exposed to different stressors such as tobacco products, smoke, particulate matters, and viruses that can activate IRE1α and may predispose individuals to bacterial infections. Studies report that certain viruses, including influenza and coronavirus, exploit activation of IRE1α signaling in epithelial cells to promote viral pathogenesis. However, whether innate immune IRE1α is needed to promote or counteract viral pathogenesis remains elusive. Our data demonstrated that the rate of body weight lost during influenza infection was the same between myeloid IRE1α-deficient mice and control littermates. Thus, this supports the idea that myeloid IRE1α does not contribute to protection against influenza infection. However, rigorous assessment of the role of myeloid IRE1α in susceptibility to different lung viral infections is needed. Viral infections are often followed by a more severe bacteria-induced pneumonia, which represents a model for a recurrent IRE1α activation in response to different stressors. To test whether repeated activation of IRE1α when exposed to different stressors contributes to inflammatory disease outcomes, we used the MRSA-induced secondary to influenza infection model. We demonstrate that loss of myeloid IRE1α is protective against influenza/MRSA coinfection, suggesting that suppression of myeloid IRE1α during repeated exposure to lung environmental stressors could be beneficial. In our future studies, we plan to broaden our findings to assess the contribution of myeloid IRE1α in disease outcomes during repeated exposure to other non-infectious lung stressors including tobacco products, smoke, particulate matter, and allergens. In summary, our data highlight the unique possibility of tuning IRE1α-mediated stress response in the lung to preprogram the protective immune response against respiratory pathogens.

## Resource availability

### Lead contact

Reagents and resources can be obtained by directing requests to the Lead Contact, Basel Abuaita.

## Acknowledgments

This work is supported by the LSU Center of Biomedical Research Excellence (COBRE) for lung biology and disease (NIH NIGMS P20GM130555-6911; B. Abuaita) and the LSU SVM startup funds to B. Abuaita. WNB is supported by the NIAID Career Transition Award (K22; AI153677). We thank the LSU Bacterial Pathogenesis and Host Response Interest Group members for many helpful discussions. We thank Drs Weishan Huang and Michael McGee for providing the influenza virus PR8. We gratefully acknowledge the services provided by the core facilities at LSU SVM including the Molecular and Cell Biology (MCBC) and Pulmonary Immunopathology (PIPC) Cores of the Center for Lung Biology and Disease (CLBD), and the Histology Core at Louisiana Animal Disease Diagnostic Laboratory (LADDL). AS was supported by Hannelore and Johannes Storz Student Travel Awards to attend national conferences.

## Author contributions

AS and BHA designed and performed the experiments; AS and BHA analyzed the data and wrote the manuscript; LMH assisted in experimental preparation and in performing the RT-qPCR experiments. WNB performed the lipidomic analysis. KH assisted in neutrophil depletion experiments. SJ assisted in experimental planning and data interpretation.

## STAR Methods

### EXPERIMENTAL MODEL AND SUBJECT DETAILS

#### Mice

Wild-type C57BL/6J and CCR2 KO mice were purchased from the Jackson Laboratory. Generation of IRE1α floxed (IRE1α^f/f^) and Myeloid IRE1α deficient (IRE1α^ΔMyeloid^) mice has been described previously.^37^ All mice were maintained according to an approved protocol in the Division of Laboratory Animal Medicine (DLAM) at the School of Veterinary Medicine, Louisiana State University and A&M College – Baton Rouge. Mice were in house genotyped using Direct mouse genotyping kit (Cat # K1025, APExBIO).

#### Bacteria

A community associated methicillin-resistant *Staphylococcus aureus* (MRSA) strain USA300 LAC was used in the study. Bacteria were maintained at –70°C in Luria-Bertani (LB) medium (Cat# 244610, Becton Dickinson) containing 20% glycerol. Bacteria were streaked onto tryptic soy medium (Cat# 211825, Becton Dickinson) plates and grown overnight at 37°C. Selected colonies were cultured overnight in tryptic soy broth (TSB) at 37°C with shaking (220 rpm). Bacteria were pelleted, washed, and resuspended in phosphate buffer saline (PBS). The bacterial inoculum was estimated based on OD_600_, and verified by plating serial dilutions onto the TSB plates to determine colony forming units (CFU).

#### Mouse lung infection

Male and female mice between the age of 8-12 weeks were used for the in *vivo* infection. Mice were anesthetized by isoflurane inhalation (3% isoflurane, 2 liter/minute oxygen). Anesthetized mice were placed on a surgical rack and inoculated with 10^7^ bacteria in 50 µl of PBS (non-lethal dose) or 10^8^ bacteria in 75 µl of PBS (lethal dose via oropharyngeal aspiration. For the influenza/MRSA coinfection, mice were inoculated with 100 plaque forming units (PFU) of influenza A virus (strain A/Puerto Rico/8/1934 H1N1, PR8) in 50 µl of PBS via oropharyngeal aspiration. PR8 was kindly provided by Dr. Huang Lab at Louisiana State University School of Veterinary Medicine. Viral infection was allowed to develop for 6 days and subsequently mice were challenged with non-lethal MRSA infection. All experimental mice were monitored at least every 6 hours for the progression of the infections. Mice were sacrificed on day 1 or 2 post-infection, BALF and lungs were collected in PBS. Lungs were mechanically dissociated into single cell suspension and further analyzed for host responses and lung damage including bacterial count enumeration, immune cell infiltration, protein and DNA accumulation in BALF, and inflammatory mediator quantification.

### METHOD DETAILS

#### Macrophage and neutrophil depletion

AM depletion was achieved by administering 50 μl of Control or Clodronate coated liposomes (Cat# CP-005-005, Liposoma BV) to mice lung via oropharyngeal aspiration in two consecutive days. The efficiency of AM depletion was determined by measuring the number of AM in the BALF on day 3 post depletion. To deplete neutrophils, mice were injected intravenously with 200 μl (0.2 mg) of rat IgG2a isotype control (Cat# BE0089, BioXCell) or an anti-mouse Ly6G monoclonal antibody (Cat# BE0075-1, BioXCell) in two consecutive days. Mice were infected with MRSA on day 3 post neutrophil depletion and the efficiency of depletion was monitored by measuring the reduction of recruited neutrophils to MRSA-infected lungs.

#### Pharmacological IRE1α inhibition *in vivo*

IRE1α small molecule inhibitor, 4μ8C (Cat# HY-19707, MedChemExpress) was dissolved in Dimethylsulfoxide (DMSO) (25 mg/ml) and then prepared for *in vivo* administration by adding the following solvent in order: 10% DMSO + 40% PEG 400 (Cat# HY-Y0873A, MedChemExpress) + 5% Tween-80 (Cat# HY-Y1891, MedChemExpress) + 45% PBS. Mice were injected 10 mg/kg of 4μ8C or vehicle control intraperitoneally 24h prior to MRSA lung infection and then every 24 hours throughout the duration of the experiment.

#### Macrophage infection

Murine alveolar macrophage cells (MH-S-CRL-2019, ATCC) were used for *in vitro* infection with MRSA. MH-S cells were cultured in Roswell Park Memorial Institute (RPMI) 1640 media containing 2 mM L-glutamine and 10% heat-inactivated fetal bovine serum (FBS). Cells were incubated at 37°C in 5% CO_2_. Macrophages were seeded in either 24-or 6-well tissue culture treated plate at a density of 2 × 10^5^ or 8 × 10^5^ cells/well respectively. The next day, cells were infected with MRSA at multiplicity of infection (MOI) of 20. Where indicated, macrophages were incubated with 4µ8C (25 mM) with Gentamicin (50 µg/ml) (Cat# 120-098-661, Quality Biological) to kill extracellular bacteria 1.5 hpi and was maintained throughout the experiment. Cells were incubated for 6 hours. Cell culture supernatants were collected and stored at –20°C for quantification of inflammatory mediators. TNF-α was analyzed by the Biolegend ELISA kit. Eicosanoids including PGE_2_, TXB_2_, LTB_4_, LTC_4_, and 15s-HETE were analyzed by Cayman Chemical ELISA kits. Cells lysates were also collected in Triton X-100 lysis buffer (50 mM Tris + 150 mM NaCl+ 1% Triton X-100) for immunoblotting analysis or in Lysis/Binding buffer (Cat# AM1561, ThermoFisher) for mRNA quantification.

#### Generation of IRE1α and Xbp1 deficient AMs

HEK293T cells were used to generate the lentivirus for CRISPR-Cas9 knockout and shRNA knockdown. HEK293T cells were grown in Dulbecco’s Modified Eagle Medium (DMEM) with 10% FBS. Cells were transfected with the TRC shRNA encoded plasmid (pLKO.1) or guided RNA (gRNA) encoded plasmid (lentiCRISPRv2) along with packaging plasmids (pHCMV-G, and pHCMV-HIV-1) using Turbofect transfection reagent (ThermoFisher). Media was changed after 24 h and virus particles were collected after 72 h post-transfection. Media-containing lentivirus was sterile filter and used to transduce macrophages. The mouse *Xbp1* specific shRNA plasmid and the non-target control shRNA plasmid (Cat# SHC016) were purchased from Millipore Sigma (Table S2). The mouse IRE1α specific gRNA and the non-target gRNA control were previously cloned into lentiCRISPRv2.^60^ Macrophages were transduced with lentivirus by mixing 1 ml of medium containing 10^6^ cells with 1 ml of medium containing virus in a 6-well plate. Cells were incubated for 48 h at 37°C in 5% CO_2_ prior selecting transduced cells with puromycin (3 μg/ml). IRE1α knockout was confirmed by the loss of protein expression using anti-IRE1α antibody (clone 14C10, Cell Signaling) or by absence of its endonuclease activity when cells were infected with MRSA. The efficiency of Xbp1 knockdown was monitored by immunoblot analysis using anti-Xbp1 antibody (clone E9V3E, Cell Signaling) when the cells were treated with ER stress inducer Thapsigargin (1 mM) (Cat# 10522, Cayman Chemical) to induce Xbp1 protein expression.

#### Immunophenotyping

BALF and lungs were collected at 48 hpi in a total of 3 ml of PBS. The lungs were mechanically dissociated into a single-cell suspension using a 70 μm strainer and red blood cells (RBC) were lysed by using an RBC lysis buffer (Cat# 420301, Biolegend). Cells were FC blocked with TruStain FcX (anti-mouse CD16/32) Antibody (Cat# 101319, Biolegend), according to the manufacturer’s instructions. Subsequently, cells were stained with the following fluorescent dye conjugated antibodies; anti-CD45 clone 30-F11, anti-CD11c clone N418, anti-CD170 clone S17007L, anti-Ly-6C clone HK1.4, anti-Ly-6G clone 1A8, anti-CD11b clone M1/70, anti-CD3 clone 17A2, anti-CD19 clone 6D5, and anti-NK-1.1 clone PK136 (Biolegend) for 30 minutes. Cells were washed and analyzed on a LSRFortessa Cell Analyzer (BD Biosciences). Compensations were performed using UltraComp Plus Compensation Beads (Cat# 01-3333, ThermoFisher). Data was further analyzed with FlowJo software. The relative abundance of immune cells in BALF and lungs was determined by quantifying the percentage of AMs (CD11c^+^, CD170^+^), neutrophils (Ly-6G^+^, CD11b^+^), monocytes (Ly6C^+^, CD11b^+^), T cells (CD3^+^), B cells (CD19^+^), and NK cells (NK1.1^+^) of total leukocytes (CD45^+^).

#### Bacterial burden and inflammatory mediator analysis

BALF and lung single cell suspensions were collected into 3 ml of PBS. BALF and lung suspensions were then diluted 1:20 in sterile ultrapure water to lyse host cells. Bacterial burden was quantified by enumerating CFU per sample after serial dilution and plating onto TSB agar plates. CFUs were counted using an Acolyte counter and confirmed by manual counting. To quantify inflammatory mediators, acellular fractions of lung single cell suspensions were collected by centrifugation and used in multiplex bead-based panels that allow quantification of multiple analytes simultaneously using Flow cytometry. The LEGENDplex mouse inflammation panel (Cat# 740446, Biolegend) and proinflammatory chemokine panel (Cat# 740451, Biolegend) were used according to the manufacturer’s protocol. Data were acquired by BD LSRFortessa flow cytometer and analyzed by LEGENDplex data analysis software suite.

#### Histology, Immunofluorescence, and TUNEL Staining

Lungs were inflated with 1.5 ml of 2% low melting point agarose in PBS and fixed with 10% neutral formalin for 2 days. The five lobes were separated and embedded in paraffin. Lungs were sectioned (5 µm per section) by the histology core at Louisiana Animal Disease Diagnostic Laboratory (LADDL) and stained with hematoxylin and eosin (H&E) staining. H&E slides were imaged on the HAMAMATSU NanoZoomer Slide Scanner System. For immunofluorescence staining, histology sections were deparaffinized and antigen retrieval was performed in sodium citrate buffer (10 mM sodium citrate, 0.05% Tween 20, pH 6.0). Sections were permeabilized with PBS + 0.2% Triton X-100 for 30 minutes and then blocked by incubating in staining buffer (PBS + 5% Bovine serum albumin (BSA, Cat# BP1600, Fisher Scientific) and 10% normal goat serum (Cat# 10000C, ThermoFisher) for 1 h. Sections were stained with the following primary antibodies overnight at 4°C: anti-T1α clone 8.1.1 (DSHB) and anti-CD169 clone 3D6.112 (Biolegend). Sections were incubated in the terminal deoxynucleotidyl transferase end labeling (TUNEL) (Cat# 11684795910, Roche) buffer for 1h at 37°C. Slides were washed with 0.1% Triton X-100 in PBS and incubated with fluorescent conjugated secondary antibodies (ThermoFisher). Sections were then counterstained with DAPI to label DNA. Slides were washed with 0.1% Triton X-100 in PBS for 20 minutes and micro cover glasses (#1 thickness Cat# 48393-106, VWR) were mounted on using ProLong Diamond Antifade Mountant (Cat# P36961, ThermoFisher). Sections were imaged using a FV3000 confocal laser scanning microscope. The percentage of TUNEL-positive cells was quantified relative to the total cells based on DAPI staining by using Image J software.

#### Cytospin and Diff-Quik staining

Cells in the BALF were immobilized onto microscope slides by centrifugation of 50 µl and 100 µl of collected BALF cells suspensions for 4 mins at 2000 rpm using Wescor Cytopro cytocentrifuge. Slides were dried overnight at room temperature and dipped sequentially into 3 steps of Quik-Dip Stain (Cat# MER 1002, Mercedes Scientific). Slides were finally dipped into clean water to wash stain solutions and dried overnight. The slides were imaged using HAMAMATSU NanoZoomer Slide Scanner System using 40X objective lens and then processed using Olympus Olyvia software.

#### Immunoblot analysis

Cell lysates were collected in Triton X-100 lysis buffer containing PhosphotaseArrest II (Cat# 786-451, G-Biosciences,) and Halt Protease Inhibitor Cocktail (Cat# 87785, ThermoFisher). Cell lysates were centrifuged, supernatant lysates were heated at 95°C for 3 min in Laemmli sample buffer (Cat# 1610747, Bio-Rad) with 10% β-mercaptoethanol (Cat# 1610710, Bio-Rad). Proteins were separated on Mini-PROTEAN TGX Stain-Free gels (Cat# 4568095, Bio-Rad) with 1X Tris/Glycine/SDS running buffer (Cat# 1610732, Bio-Rad). The gels were then transferred onto 0.45 µm nitrocellulose membranes (Cat# 10600002, Amersham) using Trans-Blot SD Semi-Dry Transfer system in 1X Tris/CAPS Buffer (Cat# 1610778, Bio-Rad). Membranes were blocked for 45 minutes in a blocking buffer (5% BSA + 0.05% Tween 20 in PBS) at room temperature shaking on an orbital shaker. Membranes were incubated with primary antibodies (1:1000 dilution) in a blocking buffer overnight at 4°C. The primary antibodies used were rabbit anti-IL-1β (D6D6T) monoclonal antibody (Cat# 31202, Cell Signaling), rabbit anti-COX 2 (D5H5) monoclonal antibody (Cat# 12282, Cell Signaling), mouse anti-GAPDH (6C5) antibody (Cat# sc-32233, Santa Cruz), and rabbit anti-β-Actin (D6A8) monoclonal antibody (Cat# 8457, Cell Signaling). Membranes were washed with PBS for at least 15 minutes and incubated with secondary LI-COR antibodies (1:5000 dilution) for 1h at room temperature. The secondary antibodies used were IRDye 800CW goat anti-mouse IgG antibody (Cat# 925-32210, LI-COR) and IRDye 680RD goat anti-rabbit IgG antibody (Cat# 925-68071, LI-COR). Membranes were washed again and scanned using the Odyssey Infrared Imaging System (LI-COR Biosciences). The images were subjected to intensity based densitometric analysis on ImageJ software to quantify the protein levels. The intensity of each band was normalized to the intensity of endogenous control (GAPDH or Actin).

#### BALF protein and DNA quantification

BALF samples collected from lungs were centrifuged and the cell free supernatants were used to quantify protein and DNA. Protein quantification was performed using Pierce BCA Protein Assay Kit (Cat# 23225, ThermoFisher) according to manufacturer’s protocol. To quantify DNA in samples, total DNA was isolated from 200 µl of normalized BALF cell free supernatants using QIAGEN DNeasy Blood & Tissue Kit (Cat# 69506, QIAGEN) and eluted in 100 µl of elution buffer. DNA levels were quantified using a NanoDrop Spectrophotometer.

#### NanoString and reverse transcription real-time quantitative PCR (RT-qPCR) analysis

Total RNA was isolated from 500 µl of lysis/binding buffer used to lyse alveolar macrophages or lung cells using mirVana miRNA isolation kit (Cat# AM1561, ThermoFisher). RNA was quantified by Nanodrop and normalized across all conditions. cDNA was synthesized using M-MLV Reverse Transcriptase (Cat# 28025013, ThermoFisher), random hexamers (Cat# N8080127, ThermoFisher), dNTP Mix (Cat# 18427088, ThermoFisher), and RNase Inhibitor (Cat# N8080119, Applied Biosystems). RT-qPCR was performed using Applied Biosystems PowerUp SYBR Green Master Mix (Cat# A25778, ThermoFisher) with the following condition: [cycle 1: 50°C for 2 mins, cycle 2: 95°C for 2 mins, cycle 3: 95°C for 15 secs, 55°C for 15 secs, 72°C for 1 min (40 repeats), cycle 4: 95°C for 15 secs, 60°C for 15 secs and 95°C 15 secs]. Endogenous controls (*Gapdh*, *Actin*) were used to calculate the relative gene expressions. The primer sequences are listed in Table S2. For NanoString assay, the nCounter mouse host response panel (NanoString Technologies) were used to monitor changes in expression of 785 inflammatory genes in MRSA infected lungs of myeloid IRE1α deficient mice and control littermates. Normalized gene counts were calculated using nSolver 4.0 software (NanoString Technologies).

#### Xbp1 splicing assay

Total RNA was extracted by using Direct-zol RNA Kit (Cat# R2052, Zymo Research) or mirVana miRNA isolation kit (Cat# AM1561, ThermoFisher). cDNA was synthesized using 500 ng of total RNA as described above. *Xbp1* transcripts (spliced and unspliced) were amplified by RT-PCR using cDNA obtained, GoTaq G2 Colorless Master Mix (Cat# M7832, Promega) and the primers listed in Table S2. Following were the PCR conditions: step 1, 95°C 2 min; step 2, 35 amplification cycles (95°C 30 s, 60°C 30 s, 72°C 30 s); and step 3, 72°C 10 min. The PCR product obtained was purified and digested with PstI restriction enzyme (Cat# R0140L, New England Biolabs) to discriminate between unspliced and spliced bands. DNA fragments were run by electrophoresis on a 2.5% agarose gel and imaged on Azure 600 Imager (Azure Biosystems).

#### Lipidomics analysis

Supernatants from lung single-cell suspensions were collected by centrifugation and stored at –20°C until analysis. Samples were thawed at 4°C for 2 h and then 500 µl of each sample was extracted twice with 800 µl of ethyl acetate. Organic/aqueous phase extraction was facilitated by centrifugation at 13,000xg for 5 min. Following centrifugation, the ethyl acetate layer was removed, combined, and dried under a stream of nitrogen gas. Each sample was dissolved in 100 µl of methanol for LC-MS/MS analysis. Following extraction, proteins in the aqueous fraction were precipitated with 1 ml of ice-cold acetone by incubating on ice for 1 h. The proteins were pelleted by centrifugation at maximum speed at 4°C for 10 min. The proteins were suspended in 200 µl of 1 % SDS in PBS, sonicated twice with a probe sonicator, and incubated for 1 h at 65°C. Proteins were quantified by BCA exactly as described in the kit instructions. LC-MS/MS analysis was performed at the LSU School of Veterinary Medicine Mass Spectrometry Resource Center on a Shimadzu 8060NX triple quadrupole mass spectrometer interfaced with a Shimadzu Nexera XS 40 series UHPLC and Shimadzu CTO-40S column oven. 5 µl of each sample, blank, or standard was analyzed in either positive or negative ionization mode, depending on the analyte. Chromatographic separation was achieved using a Phenomenex Kinetex C_8_ column (150 mm length, 2.1 mm internal diameter, 2.6 μm particle size, 100 Å pore size) with a Phenomenex Security Guard C_8_ guard column (2.1 mm internal diameter). Mobile phase A was 0.1 % formic acid in water and mobile phase B was 0.1 % formic acid in acetonitrile. The column oven was set to 40°C and the flow rate was set to 0.4 ml/min for the entire analysis. The starting condition was 10% B, which was ramped to 25% B over the first 5 min of the method. The B mobile phase was then ramped to 35 % over the next 5 min. Finally, the B mobile phase was ramped to 75% over the next 10 min. The column was washed at 98% B for 8 min, then equilibrated to 10% B for 8 min. All instrument voltages were determined and optimized empirically prior to the analysis. The Q1 *m/z*, Q3 *m/z*, retention time, and ionization mode for each analyte is listed in Table S3. The area under the curve (AUC) from each sample was normalized to total protein for differences in starting material. Fold change of normalized AUC values of infected mice relative to the average of uninfected control mice were determined for each analyte.

#### Statistical analysis

Experiments were performed at least n≥3 times on different days unless otherwise stated. Graphs represent mean ± standard deviation from biological replicates. GraphPad was used to calculate the p-values. Student’s t test was used to determine significant differences between the two groups, a One-way ANOVA with Holm-Sidak’s post-test for multiple comparisons was used to determine significant differences between three or more groups, and a Two-way ANOVA followed by Sidak’s multiple comparisons test was used to determine significant differences between groups that are defined by columns and rows. p-values of less than 0.05 were considered significant and all statistically significant comparisons within groups were considered.

## References

1. Bradley, K.L., Stokes, C.A., Marciniak, S.J., Parker, L.C., and Condliffe, A.M. (2021). Role of unfolded proteins in lung disease. Thorax 76, 92–99.

2. Schwarz, D.S., and Blower, M.D. (2016). The endoplasmic reticulum: structure, function and response to cellular signaling. Cell. Mol. Life Sci. 73, 79–94.

3. André, A.C., Laborde, M., and Marteyn, B.S. (2022). The battle for oxygen during bacterial and fungal infections. Trends Microbiol. 30, 643–653.

4. Bernatchez, J.A., and McCall, L.-I. (2020). Insights gained into respiratory infection pathogenesis using lung tissue metabolomics. PLoS Pathog. 16, e1008662.

5. Bettigole, S.E., and Glimcher, L.H. (2015). Endoplasmic reticulum stress in immunity. Annu. Rev. Immunol. 33, 107–138.

6. Walter, P., and Ron, D. (2011). The unfolded protein response: from stress pathway to homeostatic regulation. Science 334, 1081–1086.

7. Kettel, P., and Karagöz, G.E. (2024). Endoplasmic reticulum: Monitoring and maintaining protein and membrane homeostasis in the endoplasmic reticulum by the unfolded protein response. Int. J. Biochem. Cell Biol. 172, 106598.

8. van’t Wout, E.F.A., van Schadewijk, A., van Boxtel, R., Dalton, L.E., Clarke, H.J., Tommassen, J., Marciniak, S.J., and Hiemstra, P.S. (2015). Virulence Factors of Pseudomonas aeruginosa Induce Both the Unfolded Protein and Integrated Stress Responses in Airway Epithelial Cells. PLoS Pathog. 11, e1004946.

9. Oda, J.M., den Hartigh, A.B., Jackson, S.M., Tronco, A.R., and Fink, S.L. (2023). The unfolded protein response components IRE1α and XBP1 promote human coronavirus infection. MBio 14, e0054023.

10. Schmoldt, C., Vazquez-Armendariz, A.I., Shalashova, I., Selvakumar, B., Bremer, C.M., Peteranderl, C., Wasnick, R., Witte, B., Gattenlöhner, S., Fink, L., et al. (2019). IRE1 Signaling As a Putative Therapeutic Target in Influenza Virus–induced Pneumonia. Am. J. Respir. Cell Mol. Biol. 61, 537–540.

11. Cervantes-Ortiz, S.L., Zamorano Cuervo, N., and Grandvaux, N. (2016). Respiratory Syncytial Virus and Cellular Stress Responses: Impact on Replication and Physiopathology. Viruses 8. 10.3390/v8050124.

12. So, J.-S. (2018). Roles of endoplasmic reticulum stress in immune responses. Mol. Cells 41, 705–716.

13. Martinon, F., Chen, X., Lee, A.-H., and Glimcher, L.H. (2010). TLR activation of the transcription factor XBP1 regulates innate immune responses in macrophages. Nat. Immunol. 11, 411–418.

14. Tirasophon, W., Lee, K., Callaghan, B., Welihinda, A., and Kaufman, R.J. (2000). The endoribonuclease activity of mammalian IRE1 autoregulates its mRNA and is required for the unfolded protein response. Genes Dev. 14, 2725–2736.

15. Sidrauski, C., and Walter, P. (1997). The transmembrane kinase Ire1p is a site-specific endonuclease that initiates mRNA splicing in the unfolded protein response. Cell 90, 1031–1039.

16. Calfon, M., Zeng, H., Urano, F., Till, J.H., Hubbard, S.R., Harding, H.P., Clark, S.G., and Ron, D. (2002). IRE1 couples endoplasmic reticulum load to secretory capacity by processing the XBP-1 mRNA. Preprint, 10.1038/415092a 10.1038/415092a.

17. Acosta-Alvear, D., Zhou, Y., Blais, A., Tsikitis, M., Lents, N.H., Arias, C., Lennon, C.J., Kluger, Y., and Dynlacht, B.D. (2007). XBP1 Controls Diverse Cell Type– and Condition-Specific Transcriptional Regulatory Networks. Preprint, 10.1016/j.molcel.2007.06.011 10.1016/j.molcel.2007.06.011.

18. Lee, A.-H., Iwakoshi, N.N., and Glimcher, L.H. (2003). XBP-1 regulates a subset of endoplasmic reticulum resident chaperone genes in the unfolded protein response. Mol. Cell. Biol. 23, 7448–7459.

19. Hollien, J., Lin, J.H., Li, H., Stevens, N., Walter, P., and Weissman, J.S. (2009). Regulated Ire1-dependent decay of messenger RNAs in mammalian cells. J. Cell Biol. 186, 323–331.

20. So, J.-S., Hur, K.Y., Tarrio, M., Ruda, V., Frank-Kamenetsky, M., Fitzgerald, K., Koteliansky, V., Lichtman, A.H., Iwawaki, T., Glimcher, L.H., et al. (2012). Silencing of lipid metabolism genes through IRE1α-mediated mRNA decay lowers plasma lipids in mice. Cell Metab. 16, 487–499.

21. Nishitoh, H., Matsuzawa, A., Tobiume, K., Saegusa, K., Takeda, K., Inoue, K., Hori, S., Kakizuka, A., and Ichijo, H. (2002). ASK1 is essential for endoplasmic reticulum stress-induced neuronal cell death triggered by expanded polyglutamine repeats. Genes Dev. 16, 1345–1355.

22. Abuaita, B.H., Burkholder, K.M., Boles, B.R., and O’Riordan, M.X. (2015). The Endoplasmic Reticulum Stress Sensor Inositol-Requiring Enzyme 1α Augments Bacterial Killing through Sustained Oxidant Production. MBio 6, e00705.

23. Rosen, D.A., Seki, S.M., Fernández-Castañeda, A., Beiter, R.M., Eccles, J.D., Woodfolk, J.A., and Gaultier, A. (2019). Modulation of the sigma-1 receptor-IRE1 pathway is beneficial in preclinical models of inflammation and sepsis. Sci. Transl. Med. 11. 10.1126/scitranslmed.aau5266.

24. Bronner, D.N., Abuaita, B.H., Chen, X., Fitzgerald, K.A., Nuñez, G., He, Y., Yin, X.-M., and O’Riordan, M.X.D. (2015). Endoplasmic Reticulum Stress Activates the Inflammasome via NLRP3– and Caspase-2-Driven Mitochondrial Damage. Immunity 43, 451–462.

25. Abuaita, B.H., Sule, G.J., Schultz, T.L., Gao, F., Knight, J.S., and O’Riordan, M.X. (2021). The IRE1α Stress Signaling Axis Is a Key Regulator of Neutrophil Antimicrobial Effector Function. J. Immunol. 10.4049/jimmunol.2001321.

26. Turner, N.A., Sharma-Kuinkel, B.K., Maskarinec, S.A., Eichenberger, E.M., Shah, P.P., Carugati, M., Holland, T.L., and Fowler, V.G., Jr (2019). Methicillin-resistant Staphylococcus aureus: an overview of basic and clinical research. Nat. Rev. Microbiol. 17, 203–218.

27. Gastmeier, P., Kola, A., Schwab, F., Behnke, M., and Geffers, C. (2024). Etiology of nosocomial infections in intensive care patients in German hospitals: An analysis of trends between 2008 and 2022. Int. J. Med. Microbiol. 314, 151594.

28. Chopra, S., Giovanelli, P., Alvarado-Vazquez, P.A., Alonso, S., Song, M., Sandoval, T.A., Chae, C.-S., Tan, C., Fonseca, M.M., Gutierrez, S., et al. (2019). IRE1α-XBP1 signaling in leukocytes controls prostaglandin biosynthesis and pain. Science 365, eaau6499.

29. Laskin, D.L., Weinberger, B., and Laskin, J.D. (2001). Functional heterogeneity in liver and lung macrophages. J. Leukoc. Biol. 70, 163–170.

30. Vancheri, C., Mastruzzo, C., Sortino, M.A., and Crimi, N. (2004). The lung as a privileged site for the beneficial actions of PGE2. Trends Immunol. 25, 40–46.

31. Pickens, C.I., and Wunderink, R.G. (2022). Methicillin-Resistant Staphylococcus aureus Hospital-Acquired Pneumonia/Ventilator-Associated Pneumonia. Semin. Respir. Crit. Care Med. 43, 304–309.

32. Dennis, E.A., and Norris, P.C. (2015). Eicosanoid storm in infection and inflammation. Nat. Rev. Immunol. 15, 511–523.

33. Nielsen, T.B., Yan, J., Luna, B., and Spellberg, B. (2018). Murine oropharyngeal aspiration model of ventilator-associated and hospital-acquired bacterial pneumonia. J. Vis. Exp. 10.3791/57672.

34. Dobbs, L.G., Williams, M.C., and Gonzalez, R. (1988). Monoclonal antibodies specific to apical surfaces of rat alveolar type I cells bind to surfaces of cultured, but not freshly isolated, type II cells. Biochim. Biophys. Acta 970, 146–156.

35. Boring, L., Gosling, J., Chensue, S.W., Kunkel, S.L., Farese, R.V., Jr, Broxmeyer, H.E., and Charo, I.F. (1997). Impaired monocyte migration and reduced type 1 (Th1) cytokine responses in C-C chemokine receptor 2 knockout mice. J. Clin. Invest. 100, 2552–2561.

36. Iwawaki, T., Akai, R., Yamanaka, S., and Kohno, K. (2009). Function of IRE1 alpha in the placenta is essential for placental development and embryonic viability. Proc. Natl. Acad. Sci. U. S. A. 106, 16657–16662.

37. Sule, G., Abuaita, B.H., Steffes, P.A., Fernandes, A.T., Estes, S.K., Dobry, C., Pandian, D., Gudjonsson, J.E., Kahlenberg, J.M., O’Riordan, M.X., et al. (2021). Endoplasmic reticulum stress sensor IRE1α propels neutrophil hyperactivity in lupus. J. Clin. Invest. 131. 10.1172/JCI137866.

38. Heffernan, L.M., Lawrence, A.-L.E., Marcotte, H.A., Sharma, A., Jenkins, A.X., Iguwe, D., Rood, J., Herke, S.W., O’Riordan, M.X., and Abuaita, B.H. (2024). Heterogeneity of Salmonella enterica lipopolysaccharide counteracts macrophage and antimicrobial peptide defenses. Infect. Immun., e0025124.

39. Teng, F., Liu, X., Guo, S.-B., Li, Z., Ji, W.-Q., Zhang, F., and Zhu, X.-M. (2019). Community-acquired bacterial co-infection predicts severity and mortality in influenza-associated pneumonia admitted patients. J. Infect. Chemother. 25, 129–136.

40. Satjawattanavimol, S., Teerapuncharoen, K., Kaewlai, R., and Disayabutr, S. (2023). Prevalence of early bacterial co-infection in hospitalized patients with COVID-19 pneumonia: a retrospective study. J. Thorac. Dis. 15, 3568–3579.

41. Langouët-Astrié, C., Oshima, K., McMurtry, S.A., Yang, Y., Kwiecinski, J.M., LaRivière, W.B., Kavanaugh, J.S., Zakharevich, I., Hansen, K.C., Shi, D., et al. (2022). The influenza-injured lung microenvironment promotes MRSA virulence, contributing to severe secondary bacterial pneumonia. Cell Rep. 41, 111721.

42. Chertow, D.S., and Memoli, M.J. (2013). Bacterial coinfection in influenza: a grand rounds review. JAMA 309, 275–282.

43. Kitur, K., Parker, D., Nieto, P., Ahn, D.S., Cohen, T.S., Chung, S., Wachtel, S., Bueno, S., and Prince, A. (2015). Toxin-induced necroptosis is a major mechanism of Staphylococcus aureus lung damage. PLoS Pathog. 11, e1004820.

44. Zhou, Y., Niu, C., Ma, B., Xue, X., Li, Z., Chen, Z., Li, F., Zhou, S., Luo, X., and Hou, Z. (2018). Inhibiting PSMα-induced neutrophil necroptosis protects mice with MRSA pneumonia by blocking the agr system. Cell Death Dis. 9, 362.

45. Kebaier, C., Chamberland, R.R., Allen, I.C., Gao, X., Broglie, P.M., Hall, J.D., Jania, C., Doerschuk, C.M., Tilley, S.L., and Duncan, J.A. (2012). Staphylococcus aureus α-hemolysin mediates virulence in a murine model of severe pneumonia through activation of the NLRP3 inflammasome. J. Infect. Dis. 205, 807–817.

46. Cohen, T.S., Boland, M.L., Boland, B.B., Takahashi, V., Tovchigrechko, A., Lee, Y., Wilde, A.D., Mazaitis, M.J., Jones-Nelson, O., Tkaczyk, C., et al. (2018). S. Aureus evades macrophage killing through NLRP3-dependent effects on mitochondrial trafficking. Cell Rep. 22, 2431–2441.

47. Robinson, K.M., Ramanan, K., Clay, M.E., McHugh, K.J., Pilewski, M.J., Nickolich, K.L., Corey, C., Shiva, S., Wang, J., Muzumdar, R., et al. (2018). The inflammasome potentiates influenza/Staphylococcus aureus superinfection in mice. JCI Insight 3. 10.1172/jci.insight.97470.

48. Han, D., Lerner, A.G., Vande Walle, L., Upton, J.-P., Xu, W., Hagen, A., Backes, B.J., Oakes, S.A., and Papa, F.R. (2009). IRE1α kinase activation modes control alternate endoribonuclease outputs to determine divergent cell fates. J. End-to-End-test. 138, 562–575.

49. Lerner, A.G., Upton, J.-P., Praveen, P.V.K., Ghosh, R., Nakagawa, Y., Igbaria, A., Shen, S., Nguyen, V., Backes, B.J., Heiman, M., et al. (2012). IRE1α induces thioredoxin-interacting protein to activate the NLRP3 inflammasome and promote programmed cell death under irremediable ER stress. Cell Metab. 16, 250–264.

50. Estornes, Y., Aguileta, M.A., Dubuisson, C., De Keyser, J., Goossens, V., Kersse, K., Samali, A., Vandenabeele, P., and Bertrand, M.J.M. (2015). RIPK1 promotes death receptor-independent caspase-8-mediated apoptosis under unresolved ER stress conditions. Cell Death Dis. 6, e1798.

51. Greenlee-Wacker, M.C., and Nauseef, W.M. (2017). IFN-γ targets macrophage-mediated immune responses toward Staphylococcus aureus. J. Leukoc. Biol. 101, 751–758.

52. Verma, A.K., Bauer, C., Palani, S., Metzger, D.W., and Sun, K. (2021). IFN-γ drives TNF-α hyperproduction and lethal lung inflammation during antibiotic treatment of postinfluenza Staphylococcus aureus pneumonia. J. Immunol. 207, 1371–1376.

53. Verma, A.K., McKelvey, M., Uddin, M.B., Palani, S., Niu, M., Bauer, C., Shao, S., and Sun, K. (2022). IFN-γ transforms the transcriptomic landscape and triggers myeloid cell hyperresponsiveness to cause lethal lung injury. Front. Immunol. 13, 1011132.

54. Darwich, L., Coma, G., Peña, R., Bellido, R., Blanco, E.J.J., Este, J.A., Borras, F.E., Clotet, B., Ruiz, L., Rosell, A., et al. (2009). Secretion of interferon-gamma by human macrophages demonstrated at the single-cell level after costimulation with interleukin (IL)-12 plus IL-18. Immunology 126, 386–393.

55. Di Marzio, P., Puddu, P., Conti, L., Belardelli, F., and Gessani, S. (1994). Interferon gamma upregulates its own gene expression in mouse peritoneal macrophages. J. Exp. Med. 179, 1731–1736.

56. Ricciotti, E., and FitzGerald, G.A. (2011). Prostaglandins and inflammation. Arterioscler. Thromb. Vasc. Biol. 31, 986–1000.

57. Asakrah, S., Nieves, W., Mahdi, Z., Agard, M., Zea, A.H., Roy, C.J., and Morici, L.A. (2013). Post-exposure therapeutic efficacy of COX-2 inhibition against Burkholderia pseudomallei. PLoS Negl. Trop. Dis. 7, e2212.

58. Hubbard, L.L.N., Ballinger, M.N., Thomas, P.E., Wilke, C.A., Standiford, T.J., Kobayashi, K.S., Flavell, R.A., and Moore, B.B. (2010). A role for IL-1 receptor-associated kinase-M in prostaglandin E2-induced immunosuppression post-bone marrow transplantation. J. Immunol. 184, 6299–6308.

59. Lebedeva, E.S., Kuzubova, N.N., Titova, O.N., and Surkova, E.A. (2017). Effect of cyclooxygenase-2 inhibition on lung inflammation and hypoxia-inducible factor-1 signalling in COPD model. Eur. Respir. J. 50. 10.1183/1393003.congress-2017.PA3926.

60. Abuaita, B.H., Schultz, T.L., and O’Riordan, M.X. (2018). Mitochondria-Derived Vesicles Deliver Antimicrobial Reactive Oxygen Species to Control Phagosome-Localized Staphylococcus aureus. Cell Host Microbe 24, 625–636.e5.

