## Supplementary material for "Activation of the endoplasmic reticulum stress regulator IRE1α compromises pulmonary host defenses": Table S2

| Genes/Targets | Sequences |
|---|---|
| *β-actin* | Primers for RT-qPCR:<br>Forward: TGACGTTGACATCCGTAAAGACC<br>Reverse: CTCAGGAGGAGCAATGATCTTGA |
| *Gapdh* | Primers for RT-qPCR:<br>Forward: ACCACAGTCCATGCCATCAC<br>Reverse: TCCACCACCCTGTTGCTGTA |
| *Ptgs2* | Primers for RT-qPCR:<br>Forward: TGGGTGTGAAGGGAAATAAGGAG<br>Reverse: ATTTGAGCCTTGGGGGTCAG |
| *Ptges* | Primers for RT-qPCR:<br>Forward: AGCACACTGCTGGTCATCAA<br>Reverse: TTGGCAAAAGCCTTCTTCCGC |
| *Ptgs1* | Primers for RT-qPCR:<br>Forward: CTTAAGTACCAGGTGCTGGACG<br>Reverse: GGTGGGTAGCGCATCAACAC |
| *Alox15* | Primers for RT-qPCR:<br>Forward: GACACTTGGTGGCTGAGGTCTT<br>Reverse: TCTCTGAGATCAGGTCGCTCCT |
| *Alox5* | Primers for RT-qPCR:<br>Forward: TCTTCCTGGCACGACTTTGCTG<br>Reverse: GCAGCCATTCAGGAACTGGTAG |
| *Alox5ap* | Primers for RT-qPCR:<br>Forward: GTTCTTTGCCCACAAGGTGGAG<br>Reverse: TGCAGTCCAGAGTACCACAAGG |
| *Gbp2* | Primers for RT-qPCR:<br>Forward: AGATGCCCACAGAAACCCTCCA<br>Reverse: AAGGCATCTCGCTTGGCTACCA |
| *Xbp1s* | Primers for RT-qPCR:<br>Forward: GCTGAGTCCGCAGCAGGT<br>Reverse: CAGGGTCCAACTTGTCCAGAAT |
| *Xbp1* | Primers for RT-PCR:<br>Forward: AAACAGAGTAGCAGCGCAGACTGC<br>Reverse: TCCTTCTGGGTAGACCTCTGGGAG |
| Xbp1 | shRNA sense sequence:<br>GGTTGAGAACCAGGAGTTAAG |
| Non-Target Control shRNA | shRNA sequence:<br>GCGCGATAGCGCTAATAATTT |
| IRE1a | gRNA sequence:<br>CTTGTTGTTTGTCTCGACCC |
| Non-target gRNA control | gRNA sequence:<br>TCCTGCGCGATGACCGTCGG |
