## Supplementary material for "Activation of the endoplasmic reticulum stress regulator IRE1α compromises pulmonary host defenses": Table S3

| Analyte | Q1 (*m/z*) | Q3 (*m/z*) | Retention Time (min) | Ionization Mode |
|---|---|---|---|---|
| 10-HDHA | 343.2 | 153.1 | 17.2 | - |
| 10,17-DiHDHA | 359.2 | 153.1 | 14.17 | - |
| 11-dehydro-2,3-dinor-TXB2 | 339.2 | 59.1 | 8.44 | - |
| 11-dehydro-TXB2 | 367.2 | 305.2 | 10.77 | - |
| 11-dehydro-TXB3 | 365.2 | 169.1 | 9.45 | - |
| 11-HDHA | 343.2 | 149.1 | 17.29 | - |
| 11-HEDE | 323.3 | 199.1 | 18.11 | - |
| 11-HEPE | 317.2 | 121.1 | 16.225 | - |
| 11-HETE | 319.2 | 167.1 | 17.125 | - |
| 11-trans-LTC4 | 626.4 | 189.1 | 12.93 | + |
| 11-trans-LTD4 | 497.3 | 189.1 | 13 | + |
| 11-trans-LTE4 | 440.3 | 189.1 | 13.13 | + |
| 11,12-DHET | 337.2 | 167.1 | 15.36 | - |
| 11,12-EET | 319.2 | 149.1 | 18.26 | - |
| 11,12-EET-d11 | 330.3 | 167.1 | 18.2 | - |
| 11,12-EET-EA | 364.2 | 62.1 | 16.115 | + |
| 11b-13,14-dihydro-15-keto-PGF2a | 353.2 | 183.1 | 11.28 | - |
| 11b-PGE2 | 351.2 | 189.1 | 10.875 | - |
| 11b-PGF2a | 353.2 | 193.1 | 9.63 | - |
| 12-HEPE | 317.2 | 179.1 | 16.385 | - |
| 12-HETE | 319.2 | 179.1 | 17.26 | - |
| 12-HETE-d8 | 327.2 | 184.3 | 17.215 | - |
| 12-HHT | 279.2 | 179.1 | 15.155 | - |
| 12-HpEPE | 333.2 | 151.2 | 16.67 | - |
| 12-HpETE | 335.2 | 153.1 | 17.495 | - |
| 12-KETE | 317.2 | 153.1 | 17.54 | - |
| 12-keto-LTB4 | 333.2 | 179.1 | 14.74 | - |
| 12,13-DiHOME | 313.2 | 183.1 | 14.46 | - |
| 12,13-EpOME | 295.2 | 195.2 | 17.76 | - |
| 12(13) - EPOME-d4 | 299.5 | 198.5 | 17.76 | - |
| 12(13)- DiHOME-d4 | 317.5 | 185.5 | 14.46 | - |
| 13 - HODE - d4 | 299.5 | 198.5 | 16.6 | - |
| 13 - KODE-d3 | 296.5 | 198.5 | 17 | - |
| 13-HDHA | 343.2 | 193.1 | 17.15 | - |
| 13-HODE | 295.2 | 195.1 | 16.6 | - |
| 13-HOTrE | 293.2 | 195.1 | 15.85 | - |
| 13-HpODE | 311.2 | 113.1 | 17.01 | - |
| 13-HpOTrE | 309.2 | 195.1 | 16.26 | - |
| 13-KODE | 293.2 | 113.1 | 17 | - |
| 13,14-dihydro-15-keto PGJ2 | 333.2 | 113 | 14.18 | - |
| 13,14-dihydro-15-keto-PGA2 | 333.2 | 175.1 | 14.005 | - |

| | | | | |
|---|---|---|---|---|
| 13,14-dihydro-15-keto-PGD1 | 353.2 | 221.1 | 12.935 | - |
| 13,14-dihydro-15-keto-PGD2 | 351.2 | 207.2 | 12.54 | - |
| 13,14-dihydro-15-keto-PGE2 | 351.2 | 175.1 | 11.895 | - |
| 13,14-dihydro-15-keto-PGF2a | 353.2 | 113.1 | 11.72 | - |
| 13,14-dihydro-15-keto-tetranor-PGD2 | 297.2 | 109.1 | 7.9 | - |
| 13,14-dihydro-15-keto-tetranor-PGE2 | 297.2 | 109.1 | 8.59 | - |
| 13,14-dihydro-15-keto-tetranor-PGF1a | 299.2 | 113.1 | 7.52 | - |
| 13,14-dihydro-15-keto-tetranor-PGF1b | 299.2 | 113.1 | 6.62 | - |
| 13,14-dihydro-PGE1 | 355.2 | 193.1 | 11.595 | - |
| 13,14-dihydro-PGF1a | 357.2 | 113.4 | 11.5 | - |
| 14-HDHA | 343.2 | 205.2 | 17.24 | - |
| 14,15-DHET | 337.2 | 207.2 | 15.04 | - |
| 14,15-DiHET-d11 | 348.3 | 207.1 | 14.978 | - |
| 14,15-DiHETE | 335.2 | 207.2 | 14.31 | - |
| 14,15-EET | 319.2 | 113.1 | 17.95 | - |
| 14,15-EET-EA | 364.2 | 62.1 | 15.75 | + |
| 14,15-EpETE | 317.2 | 207.2 | 17.28 | - |
| 14,15-LTC4 | 626.4 | 308.2 | 11.67 | + |
| 14,15-LTE4 | 440.3 | 301.2 | 12.24 | + |
| 15-deoxy-delta-12,14-PGJ2 | 315.2 | 271.2 | 15.97 | - |
| 15-HEDE | 323.2 | 223.2 | 18.155 | - |
| 15-HEPE | 317.2 | 219.2 | 16.2 | - |
| 15-HETE | 319.2 | 219.2 | 16.93 | - |
| 15-HETE-d8 | 327.2 | 226.2 | 16.86 | - |
| 15-HETrE | 321.2 | 221.2 | 17.42 | - |
| 15-HpEPE | 333.2 | 111.1 | 16.56 | - |
| 15-HpETE | 335.2 | 113.1 | 17.27 | - |
| 15-KEDE | 323.2 | 207.2 | 18.54 | + |
| 15-KETE | 317.2 | 113.1 | 17.265 | - |
| 15-keto-PGE2 | 349.2 | 113.1 | 11.285 | - |
| 15-keto-PGF1a | 353.2 | 193.1 | 11.205 | - |
| 15-keto-PGF2a | 351.2 | 219.2 | 10.815 | - |
| 16-HDHA | 343.2 | 233.2 | 17.05 | - |
| 16-HETE | 319.2 | 233.2 | 16.445 | - |
| 16,17-EpDPA | 343.2 | 147.1 | 18.15 | - |
| 17-HDHA | 343.2 | 245.2 | 17.08 | - |
| 17-HETE | 319.2 | 247.2 | 16.36 | - |
| 17-HpDHA | 359.2 | 111.1 | 17.39 | - |
| 17,18-DiHETE | 335.2 | 247.2 | 13.94 | - |
| 17,18-EpETE | 317.2 | 255.2 | 16.96 | - |
| 18-carboxy-dinor-LTB4 | 337.2 | 59.1 | 6.09 | - |
| 18-HEPE | 317.2 | 215.2 | 15.86 | - |
| 18-HETE | 319.2 | 261.2 | 16.245 | - |

| | | | | |
|---|---|---|---|---|
| 19-HETE | 319.2 | 275.2 | 15.95 | - |
| 19,20-DiHDPA | 361.2 | 87.1 | 15.01 | - |
| 19,20-EpDPA | 343.2 | 241.2 | 17.835 | - |
| 1a,1b-dihomo-PGF2a | 381.2 | 337.2 | 12.37 | - |
| 2,3-dinor-11b-PGF2a | 325.2 | 145.1 | 7.6 | - |
| 2,3-dinor-8-iso-PGF2a | 325.2 | 237.2 | 7.22 | - |
| 2,3-dinor-PGE1 | 325.2 | 245.2 | 8.625 | - |
| 2,3-dinor-TXB1 | 343.2 | 143.1 | 6.985 | - |
| 2,3-dinor-TXB2 | 341.2 | 123.1 | 7.36 | - |
| 20-carboxy-AA | 333.2 | 297.2 | 15.64 | - |
| 20-carboxy-LTB4 | 365.2 | 169.1 | 8.05 | - |
| 20-HDHA | 343.2 | 241.2 | 16.77 | - |
| 20-HETE | 319.2 | 245.3 | 16.07 | - |
| 20-hydroxy-LTB4 | 351.2 | 195.1 | 8.34 | - |
| 20-hydroxy-PGE2 | 367.2 | 287.2 | 5.58 | - |
| 20-hydroxy-PGF2a or 19-hydroxy-PGF2a | 369.2 | 325.2 | 5.43 | - |
| 4-HDHA | 343.2 | 101.1 | 17.65 | - |
| 5-HEPE | 317.2 | 115.1 | 16.45 | - |
| 5-HETE | 319.2 | 115.1 | 17.36 | - |
| 5-HETE-d8 | 327.2 | 116.1 | 17.3 | - |
| 5-HETrE | 321.2 | 115.1 | 18.295 | - |
| 5-HpEPE | 333.2 | 173.1 | 16.865 | - |
| 5-HpETE | 335.2 | 129.1 | 17.73 | - |
| 5-iPF2a-VI | 353.2 | 115.1 | 9.78 | - |
| 5-KETE | 317.2 | 203.2 | 18.04 | - |
| 5,15-DiHETE | 335.2 | 173.1 | 14 | - |
| 5,6-DHET | 337.2 | 145.1 | 15.8 | - |
| 5,6-DHET-lactone | 321.2 | 177.1 | 17.62 | + |
| 5,6-DiHETE | 335.2 | 145.1 | 14.84 | - |
| 5,6-EET | 319.2 | 191.1 | 18.34 | - |
| 5,6-EET-EA | 364.2 | 62.1 | 16.44 | + |
| 5S,14R-LXB4 | 351.3 | 221.2 | 10.98 | - |
| 5S,6R-LXA4 | 351.2 | 115.1 | 11.805 | - |
| 5S,6S-LXA4 | 351.2 | 115.1 | 12.06 | - |
| 6-keto-PGE1 | 367.2 | 143.1 | 8.11 | - |
| 6-keto-PGF1a | 369.2 | 245.2 | 7.68 | - |
| 6-keto-PGF1a-d4 | 373.2 | 249.1 | 7.64 | - |
| 6-trans-LTB4 | 335.2 | 195.1 | 14 | - |
| 6,15-diketo-13,14-dihydro-PGF1a | 369.2 | 113.1 | 8.87 | - |
| 7-HDHA | 343.2 | 141.1 | 17.29 | - |
| 7,17-DiHDPA | 361.2 | 143.1 | 14.41 | - |
| 8-HDHA | 343.2 | 109.1 | 17.35 | - |
| 8-HEPE | 317.2 | 127.1 | 16.25 | - |

| | | | | |
|---|---|---|---|---|
| 8-HETE | 319.2 | 155.1 | 17.2 | - |
| 8-HETrE | 321.2 | 157.2 | 17.49 | - |
| 8-iso-13,14-dihydro-15-keto-PGF2a | 353.2 | 183.1 | 10.46 | - |
| 8-iso-15-keto-PGF2a | 351.2 | 219.2 | 9.94 | - |
| 8-iso-15R-PGF2a | 353.2 | 193.1 | 9.27 | - |
| 8-iso-PGA1 | 335.2 | 235.2 | 13.035 | - |
| 8-iso-PGA2 | 333.2 | 271.2 | 12.92 | - |
| 8-iso-PGE1 | 353.2 | 235.2 | 10.795 | - |
| 8-iso-PGE2 | 351.2 | 271.2 | 10.54 | - |
| 8-iso-PGF1a | 355.2 | 293.2 | 9.485 | - |
| 8-iso-PGF2a | 353.2 | 193.1 | 9.425 | - |
| 8-iso-PGF3a | 351.2 | 307.2 | 8.24 | - |
| 8,12-iso-iPF2a-VI-1,5-lactone | 337.2 | 265.2 | 13.8 | + |
| 8,15-DiHETE | 335.2 | 127.1 | 13.81 | - |
| 8,9-DHET | 337.2 | 127.1 | 15.54 | - |
| 8,9-EET | 319.2 | 127.1 | 18.33 | - |
| 8,9-EET-EA | 364.2 | 62.1 | 16.26 | + |
| 9-HEPE | 317.2 | 69.35 | 16.355 | - |
| 9-HETE | 319.2 | 123.1 | 17.295 | - |
| 9-HODE | 295.2 | 171.1 | 16.6 | - |
| 9-HOTrE | 293.2 | 171.1 | 15.68 | - |
| 9-HpODE | 311.2 | 185.2 | 16.995 | - |
| 9-KODE | 293.2 | 185.1 | 17.12 | - |
| 9,10-DiHOME | 313.2 | 201.2 | 14.63 | - |
| 9,10-EpOME | 295.2 | 171.1 | 17.83 | - |
| Azelaoyl-PAF | 650.4 | 201.2 | 18.1 | - |
| delta17-6-keto-PGF1a | 367.2 | 163.1 | 6.57 | - |
| iPF2a-IV | 353.2 | 127.1 | 9.07 | - |
| LTB3 | 337.2 | 123 | 14.98 | - |
| LTB4 | 335.2 | 195.1 | 14.12 | - |
| LTB4 d4 | 339.2 | 197.1 | 14.095 | - |
| LTB4-EA | 362.2 | 189.1 | 12.57 | + |
| LTB5 | 333.2 | 195.1 | 13.14 | - |
| LTC4 | 626.4 | 308.2 | 12.62 | + |
| LTC4-d5 | 631.4 | 308.2 | 12.59 | + |
| LTD4 | 497.3 | 189.1 | 12.67 | + |
| LTD4-d5 | 502.3 | 194.2 | 12.65 | + |
| LTE4 | 440.3 | 189.1 | 12.86 | + |
| LTF4 | 569.4 | 251.2 | 12.87 | + |
| LXA5 | 349.2 | 115.1 | 10.47 | - |
| Lyso-PAF | 482.3 | 104.2 | 16.49 | + |
| Maresin1 | 359.2 | 113.1 | 14.09 | - |
| N-acetyl-LTE4 | 480.3 | 333.2 | 14.905 | - |

| | | | | |
|---|---|---|---|---|
| PAF | 568.4 | 59.1 | 17.5 | - |
| PAF-d4 | 572.4 | 59.1 | 17.485 | - |
| PGA1 | 335.2 | 235.2 | 13.375 | - |
| PGA2 | 333.2 | 189.1 | 13.065 | - |
| PGA2-d4 | 337.2 | 275.2 | 13.04 | - |
| PGB2 | 333.2 | 175.1 | 13.195 | - |
| PGD1 | 353.2 | 235.1 | 11.205 | - |
| PGD2 | 351.2 | 271.2 | 11.1 | - |
| PGD2-d4 | 355.2 | 275.2 | 11.08 | - |
| PGD2-EA | 440.3 | 271.2 | 8.64 | - |
| PGD3 | 349.2 | 269.2 | 9.8 | - |
| PGE1 | 353.2 | 235.2 | 11.08 | - |
| PGE1-EA | 442.3 | 360.2 | 8.41 | - |
| PGE2 | 351.2 | 271.2 | 10.7 | - |
| PGE2-d4 | 355.2 | 275.2 | 10.67 | - |
| PGE2-EA | 440.3 | 271.2 | 8.2 | - |
| PGE3 | 349.2 | 269.2 | 9.47 | - |
| PGF1a | 355.2 | 211.1 | 10.44 | - |
| PGF2a | 353.2 | 193.1 | 10.27 | - |
| PGF2a-d4 | 357.2 | 197.2 | 10.25 | - |
| PGF2a-EA | 442.3 | 334.2 | 8.065 | - |
| PGF3a | 351.2 | 193.1 | 9.04 | - |
| PGJ2 | 333.2 | 233.2 | 13.17 | - |
| PGK2 | 349.2 | 249.2 | 11.07 | - |
| Resolvin D1 | 375.2 | 141.1 | 11.96 | - |
| Resolvin D2 | 375.2 | 175.1 | 11.355 | - |
| Resolvin D3 | 375.2 | 147.1 | 10.885 | - |
| Resolvin D4 | 375.2 | 131.1 | 12.83 | - |
| Resolvin D5 | 359.2 | 199.1 | 14.215 | - |
| Resolvin E1 | 349.2 | 161.1 | 7.96 | - |
| tetranor-12-HETE | 265.2 | 109.1 | 14.85 | - |
| tetranor-PGAM | 309.1 | 163.1 | 4.585 | - |
| tetranor-PGDM | 327.2 | 309.2 | 3.26 | - |
| tetranor-PGEM | 327.2 | 309.2 | 3.02 | - |
| tetranor-PGEM-d6 | 333.2 | 315.2 | 2.99 | - |
| tetranor-PGFM | 329.2 | 311.2 | 2.66 | - |
| tetranor-PGJM | 309.1 | 155.1 | 4.46 | - |
| TXB1 | 371.2 | 171.2 | 9.285 | - |
| TXB2 | 369.2 | 195.1 | 9.465 | - |
| TXB2-d4 | 373.2 | 199.1 | 9.44 | - |
| TXB3 | 367.2 | 169.1 | 8.275 | - |
