## Supplemental Figures 1-5 for "Activation of the endoplasmic reticulum stress regulator IRE1α compromises pulmonary host defenses"

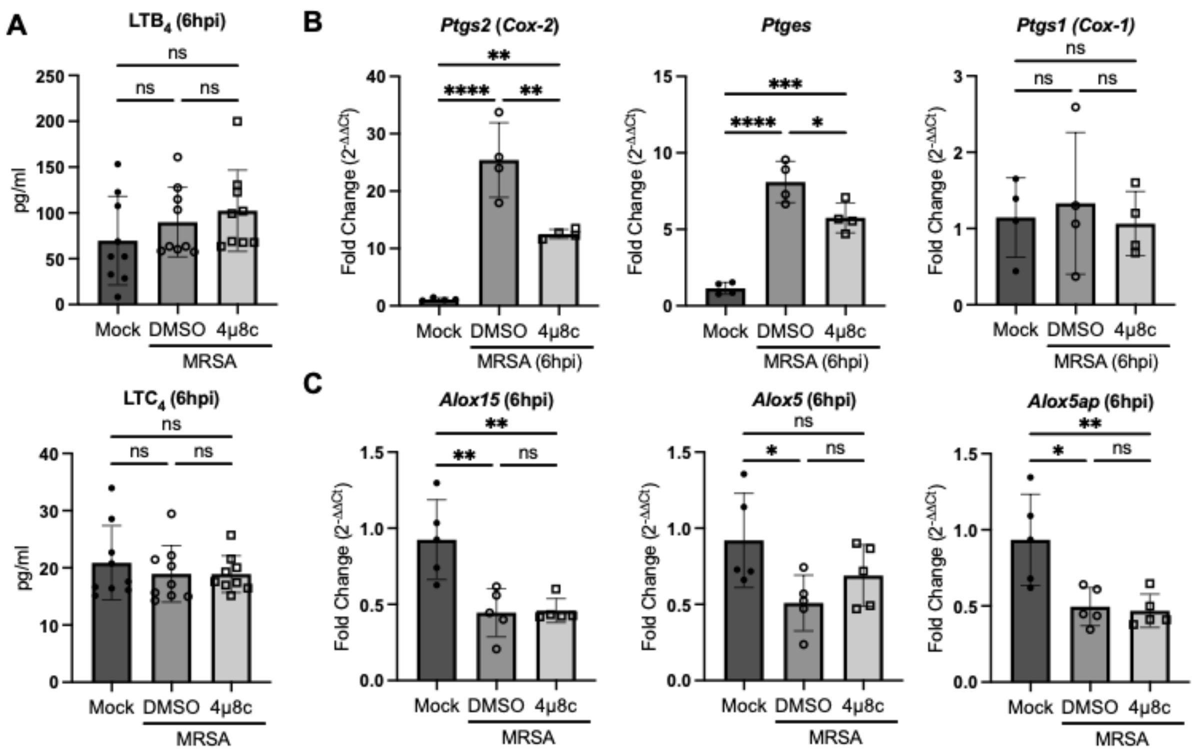


**Supplemental Figure 1: IRE1α selectively promotes expression enzymes of cyclooxygenase pathway during MRSA infection. (A)** Levels of $LTB_4$ and $LTC_4$ in culture media of AMs when cells were left untreated (Mock) or infected with MRSA for 6 h in the presence and absence of 4μ8C. Eicosanoids were quantified by ELISA. **(B, C)** Quantitative RT-qPCR of *Ptgs2*, *Ptgs1*, *Alox15*, *Alox5* and *Alox5ap* transcript levels of AMs when cells were left untreated (Mock) or infected with MRSA for 6 h in the presence and absence of 4μ8C. Graphs indicate the mean of n≥3 independent experiments ± SD. Statistical analysis was calculated by One-way ANOVA with Holm-Sidak's post-test for multiple comparisons. *$P < 0.05$; **$P < 0.01$, ***$P < 0.001$, ****$P < 0.0001$.

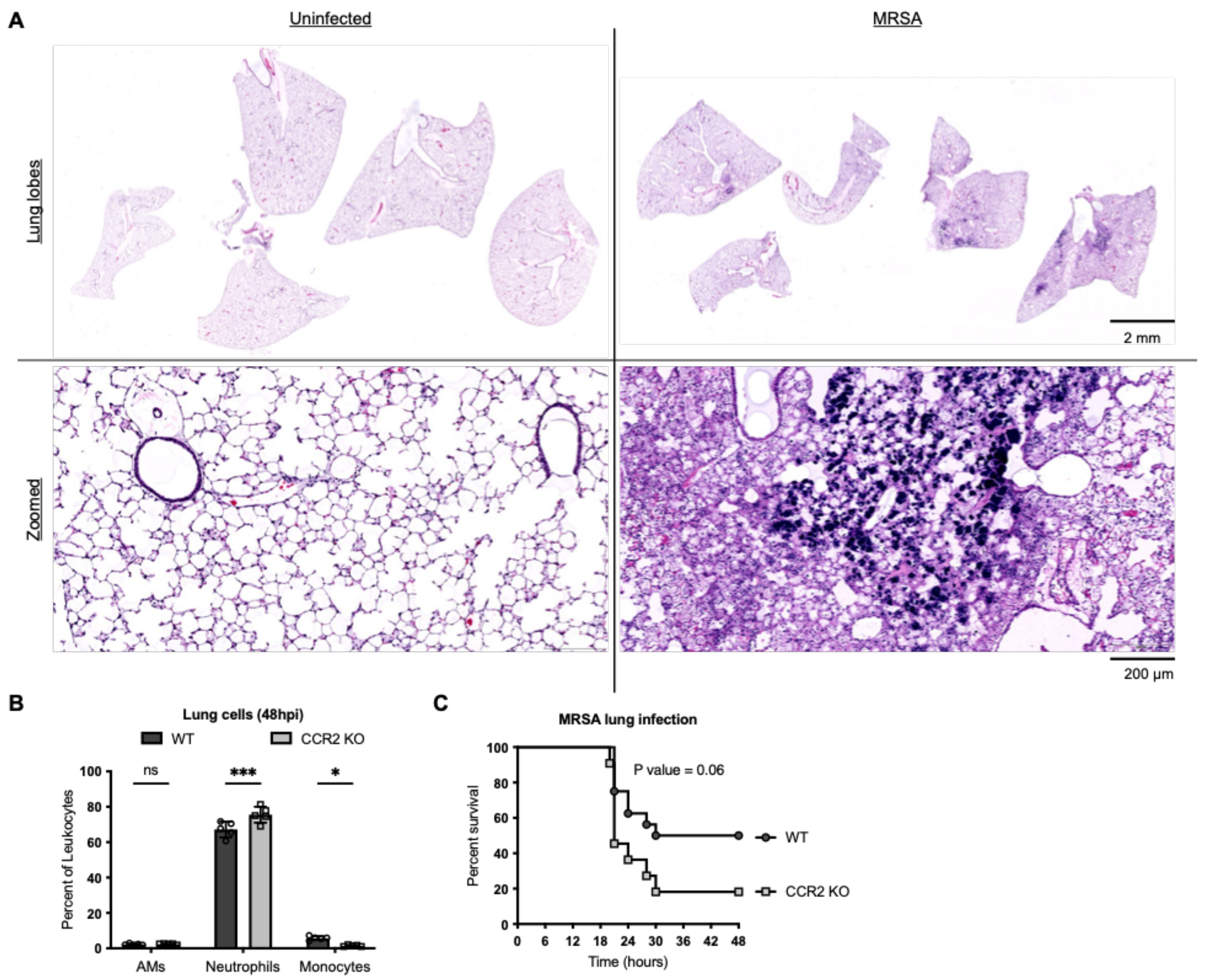


**Supplemental Figure 2: Monocytes are required for immunity against MRSA-induced pneumonia.** (A) Hematoxylin and eosin (H&E) stained lung histology sections from uninfected or MRSA infected mice for 48 h. (B) Immunophenotyping of leukocytes in the lungs of WT and CCR2 KO when infected with MRSA via oropharyngeal aspiration for 48 h. Cells were stained for AM markers (anti-CD11c, anti-CD170), neutrophil (PMNs) markers (anti-Ly6G, anti-CD11b), and monocyte (MONO) markers (anti Ly6C, anti-CD11b) and analyzed by flow cytometry. Graph indicates mean of n=5 WT and n=5 CCR2 KO mice from two independent experiments. p-values were calculated using Two-way ANOVA with Sidak's post-test. P value: *< 0.05 and *** < 0.001. (C) Survival curve of

MRSA infected WT and CCR2 KO mice via oropharyngeal aspiration for 48 h. P Value was calculated by the Log-rank (Mantel-Cox) test.

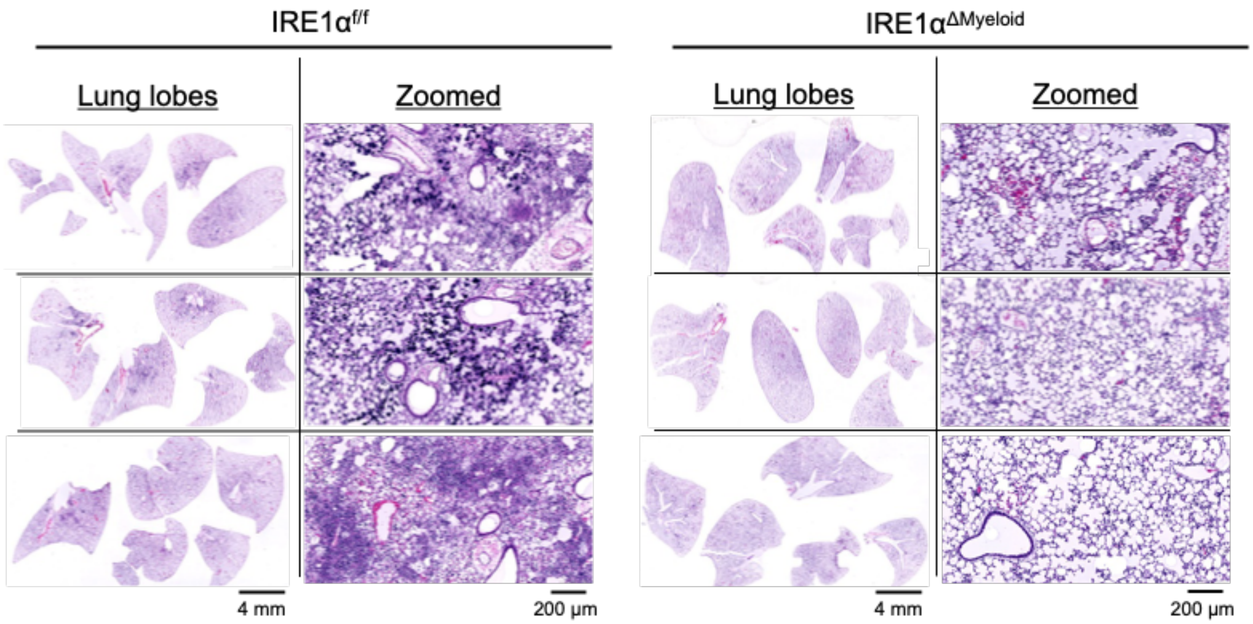


**Supplemental Figure 3: Myeloid IRE1α promotes MRSA-induced lung injury.** Representative images of H&E-stained lung histology sections from myeloid IRE1α deficient (IRE1α$^{\Delta Myeloid}$) and control littermate (IRE1α$^{f/f}$) mice when infected with MRSA via oropharyngeal route for 24 hours.

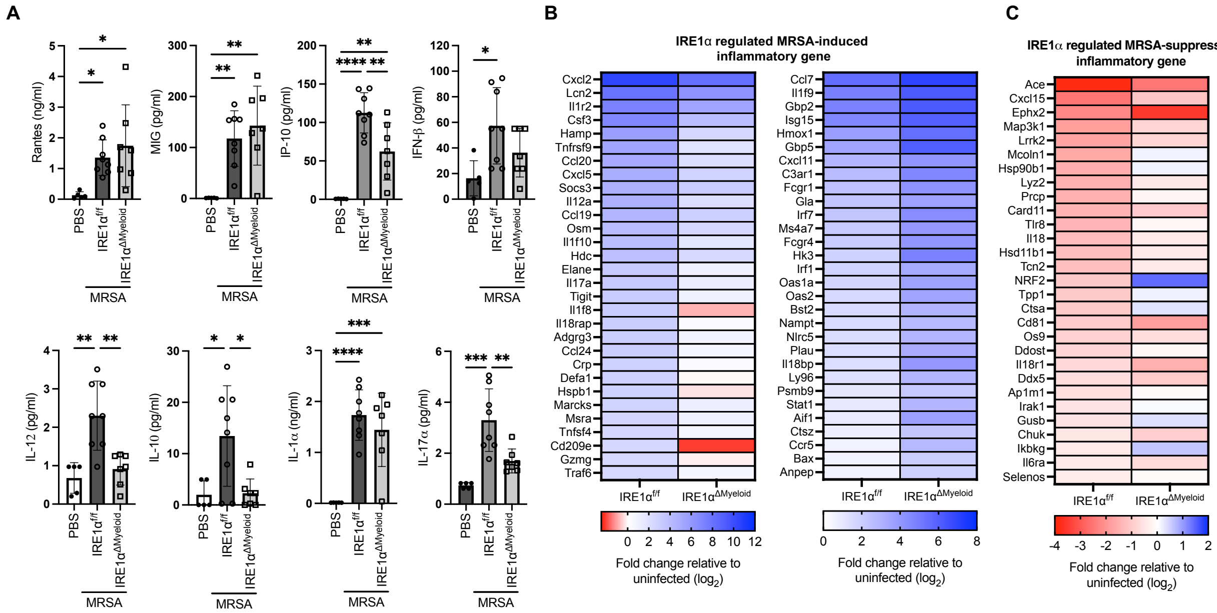


**Supplemental Figure 4: Silencing myeloid IRE1α interferes with expression of several inflammatory genes in response to MRSA lung infection.** (A) Cytokine and chemokine levels in lungs of uninfected, MRSA-infected IRE1α$^{f/f}$, and MRSA-infected IRE1α$^{\Delta Myeloid}$ mice were analyzed by flow cytometry using a bead-based multiplex assay. (B) Heatmaps depicting IRE1α-dependent positively (left) and negatively (right) regulated genes based on significant genes sorted by the greatest difference in upregulated genes between MRSA-infected IRE1α$^{f/f}$ relative to uninfected mice and MRSA infected IRE1α$^{\Delta Myeloid}$ relative to uninfected mice at 48 hpi. (C) Heatmap depicting IRE1α-dependent suppressed genes in response to MRSA. Genes were sorted based on the greatest difference between the downregulated genes in MRSA infected IRE1α$^{f/f}$ relative to uninfected mice and MRSA infected IRE1α$^{\Delta Myeloid}$ relative to uninfected mice at 48 hpi. Graphs representing the mean of n≥5 biological replicates ± SD. Statistical analysis was performed by One-way ANOVA with Holm-Sidak's post-test for multiple comparisons. *$P < 0.05$; **$P < 0.01$, ***$P < 0.001$, ****$P < 0.0001$.

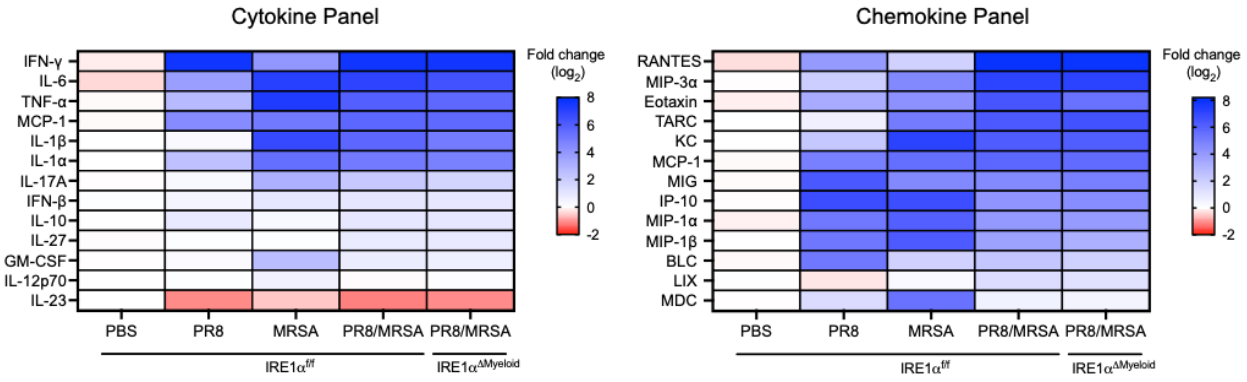


**Supplemental Figure 5: Production of cytokine and chemokine in lungs of uninfected (PBS), influenza PR8 infected, and influenza PR8/MRSA coinfected mice.** Supernatants from single-cell lung suspensions of indicated groups were used to analyze the levels of cytokines and chemokines by flow cytometry using a bead-based multiplex assay. Cytokine and chemokine levels presented as $\log_2$ (fold change) relative to uninfected (PBS) mice. Heatmaps present data from n≥4 mice of each group.
