## Supplementary material for "Activation of the endoplasmic reticulum stress regulator IRE1α compromises pulmonary host defenses": Table S1

| Probe Name | Fold change ($log_{10}$) | Significance ($-Log_{10}pValue$) |
|---|---|---|
| Ager | 0.673715794 | 6.25255748 |
| Mcoln1 | -0.540968848 | 5.27687706 |
| Tnfrsf9 | 1.046691073 | 5.1918593 |
| Hsp90b1 | -0.514199591 | 5.16972487 |
| Psap | -0.780049254 | 5.13760492 |
| Strap | 0.313900161 | 5.13573738 |
| Prcp | -0.392747975 | 4.83592663 |
| Il16 | 0.412699767 | 4.81991306 |
| Stat5a | 0.378436384 | 4.76548376 |
| Mapk13 | 0.769102125 | 4.58722031 |
| Cd68 | -0.711014255 | 4.57852468 |
| Tpp1 | -0.367702303 | 4.48763411 |
| Cdh1 | 0.491536633 | 4.44097136 |
| Ccl27a/b | 1.333356312 | 4.34047206 |
| Hamp | 0.94490353 | 4.11148561 |
| Notch1 | 0.335352405 | 4.10394553 |
| Cklf | -0.514739691 | 3.95787948 |
| Myc | 0.630643758 | 3.78121822 |
| Il18rap | 0.825897684 | 3.74805704 |
| Hspb1 | 0.922922656 | 3.66248908 |
| Map3k3 | 0.313341606 | 3.63162954 |
| Gstm4 | 0.823673272 | 3.59518241 |
| NRF2 | -0.742380809 | 3.59145586 |
| Map2k4 | 0.224600793 | 3.51669419 |
| Gzmf | 0.707661643 | 3.50654905 |
| Stk11ip | 0.234482965 | 3.47333463 |
| Il19 | 0.874269477 | 3.46246469 |
| Ccl20 | 0.905658089 | 3.44209679 |
| Ctsa | -0.407900229 | 3.4376047 |
| Cystm1 | 0.689050881 | 3.41208093 |
| Tnf | -0.355710315 | 3.38312719 |
| Nt5e | 0.835416256 | 3.38090637 |
| Plscr1 | 0.667460724 | 3.37573534 |
| Ptger2 | -0.553348869 | 3.37440589 |
| Bax | -0.431712097 | 3.33519984 |
| Atg7 | -0.549130737 | 3.22469813 |
| Ep300 | 0.13898104 | 3.21467977 |
| Igfbp7 | 0.223558511 | 3.18644064 |
| Gzmg | 0.767061148 | 3.1323934 |
| Wipi1 | 0.619115138 | 3.10616693 |

| | | |
|---|---|---|
| Oas2 | -0.588253032 | 3.10412404 |
| Il1f5 | 1.30596732 | 3.07888389 |
| Itgal | -0.370401991 | 3.06327783 |
| Map2k3 | 0.552676938 | 3.0576884 |
| Gba | -0.332863133 | 3.02647758 |
| Tollip | 0.142286707 | 3.02191894 |
| Map3k8 | 0.439128414 | 3.01010857 |
| Ms4a2 | 0.949719172 | 3.00817201 |
| Akt1 | 0.109782468 | 2.97888569 |
| Plcg2 | -0.236610773 | 2.97018367 |
| Bcl6 | -0.372432776 | 2.97014438 |
| Map3k1 | -0.311188116 | 2.96382136 |
| Tgfbr2 | 0.376370777 | 2.95776291 |
| Map1lc3a | 0.395528093 | 2.94208917 |
| Il17ra | 0.264970126 | 2.92845842 |
| Eif3f | 0.227411962 | 2.92582937 |
| Tcn2 | -0.233119895 | 2.91976533 |
| Ctsz | -0.469374703 | 2.91805384 |
| Socs3 | 0.469544858 | 2.81951496 |
| Ikbkg | -0.307777182 | 2.79872527 |
| Mapk8 | 0.427159675 | 2.75470547 |
| Ly96 | -0.493664975 | 2.74436502 |
| Cd84 | -0.410229932 | 2.73862824 |
| Traf6 | 0.562859816 | 2.73120692 |
| Cxcl14 | 0.379950652 | 2.71106733 |
| Ifitm2 | 0.432431837 | 2.7048627 |
| Vamp3 | 0.246536568 | 2.68380711 |
| Fas | 0.224513492 | 2.66303473 |
| Ccr5 | -0.662079287 | 2.65644767 |
| Rel | 0.416597836 | 2.65273165 |
| H2-Q2 | 0.454025863 | 2.6464044 |
| Il1f9 | -0.393963567 | 2.63483504 |
| Csf1 | 0.395075246 | 2.63162894 |
| Atp6v1b2 | -0.28642451 | 2.61599213 |
| Kras | 0.359257271 | 2.60732691 |
| Cd209e | 1.363168795 | 2.60311518 |
| Alox5 | 0.525908543 | 2.58632788 |
| Il6ra | 0.092691591 | 2.58141797 |
| Mapkapk2 | 0.25555767 | 2.57253424 |
| F5 | 0.860241444 | 2.55947609 |
| Txnip | 0.415605263 | 2.55839688 |

| | | |
|---|---|---|
| Relb | 0.221703942 | 2.54731 |
| Il2rg | -0.558530572 | 2.54443932 |
| Tmprss2 | 0.786735674 | 2.5321426 |
| Gab2 | 0.270861624 | 2.52709845 |
| Nfkbia | 0.311056116 | 2.51911252 |
| Mrps7 | 0.356004432 | 2.50745871 |
| Havcr2 | 0.268794258 | 2.49112967 |
| Ccr2 | -0.42706954 | 2.49048793 |
| Gusb | -0.25967702 | 2.4762495 |
| Hk3 | -0.884082301 | 2.46615186 |
| Ifi203 | -0.611099779 | 2.46528881 |
| Lif | 0.957526646 | 2.46186485 |
| Elane | 0.69499135 | 2.45947775 |
| Ube2n | 0.196860491 | 2.45895118 |
| Hcst | 0.468553634 | 2.4236222 |
| Ccr1l1 | 0.844381855 | 2.40362487 |
| Cd27 | 1.171950266 | 2.40083101 |
| H2-Q10 | 0.380505389 | 2.39791793 |
| Cxcl5 | 0.627976923 | 2.3922086 |
| Lamp1 | -0.269457613 | 2.38455335 |
| Il10rb | -0.271934104 | 2.38338759 |
| Scarb2 | 0.142102195 | 2.37581456 |
| Il5ra | 0.604804835 | 2.35320765 |
| Pdhb | 0.302277821 | 2.35131516 |
| Ahr | 0.414661788 | 2.349974 |
| Adgrg3 | 0.681621135 | 2.3380726 |
| Bcr | 0.664496659 | 2.32053851 |
| Pik3r4 | 0.322591086 | 2.30794689 |
| Oas1a | -0.58861923 | 2.3072218 |
| Gca | -0.284445138 | 2.30429225 |
| Cd44 | 0.343283336 | 2.301954 |
| Cyp2e1 | 0.718896886 | 2.28419157 |
| Irak1 | -0.189037981 | 2.27360231 |
| Apex1 | 0.400084813 | 2.26104129 |
| Ywhaq | 0.295086845 | 2.25867994 |
| Gzmb | 0.585988563 | 2.25833087 |
| Icosl | 0.250592989 | 2.25705657 |
| Il1f8 | 1.196030638 | 2.24438188 |
| Stat1 | -0.58191485 | 2.24310141 |
| Csf2 | 1.047118173 | 2.24016108 |
| Foxo1 | 0.295937683 | 2.23039178 |

| | | |
|---|---|---|
| Pik3cg | 0.192813116 | 2.22688874 |
| Adora2a | 0.292482086 | 2.22686132 |
| Rb1cc1 | 0.398545324 | 2.2191993 |
| Ackr4 | 0.614947541 | 2.21312231 |
| Atp6ap2 | 0.207488413 | 2.21232451 |
| Fos | 0.394841229 | 2.2082499 |
| Ccl24 | 0.732456014 | 2.19375012 |
| Ntng2 | 0.339305372 | 2.1866811 |
| Defa1 | 0.87253323 | 2.18262474 |
| Crk | 0.301919446 | 2.17606956 |
| Serpina1a | 0.746769794 | 2.17554657 |
| Trim6 | 0.644453915 | 2.17516429 |
| Il4ra | 0.308295768 | 2.16199061 |
| Becn1 | 0.186965799 | 2.15635578 |
| Tlr3 | 0.632231308 | 2.15280345 |
| Gla | -0.385765871 | 2.14852945 |
| Irak4 | -0.252883291 | 2.14565711 |
| Il12a | 0.78910535 | 2.14560739 |
| Cxcr5 | 0.685598232 | 2.14527026 |
| Nox1 | 0.620736164 | 2.14480527 |
| Sirpa | -0.263142923 | 2.14140579 |
| Peli2 | 0.326031793 | 2.14043938 |
| Hmox1 | -0.507476003 | 2.13423874 |
| Mmp9 | 0.334161838 | 2.13267032 |
| Rbpj | 0.143728943 | 2.11185627 |
| C1qbp | 0.19475695 | 2.11089489 |
| Ctss | -0.563575536 | 2.11014402 |
| Stat4 | 0.554299397 | 2.10110713 |
| Sod1 | 0.339880593 | 2.0851068 |
| Crp | 0.790161272 | 2.08067334 |
| Junb | 0.35836217 | 2.0802543 |
| Klrc1 | 0.521751134 | 2.05971583 |
| Nlrc5 | -0.443021308 | 2.05473926 |
| Prdm1 | 0.545252906 | 2.05188704 |
| Tank | -0.118199119 | 2.04472714 |
| Atp6v0d1 | -0.243193032 | 2.04229774 |
| Tigit | 0.652177098 | 2.03888025 |
| Plin4 | 0.731445093 | 2.03319644 |
| Itm2c | 0.265395978 | 2.02485882 |
| Gbp5 | -0.758645714 | 2.02279964 |
| Ulk1 | 0.412665443 | 2.00256261 |

| | | |
|---|---|---|
| Trim56 | -0.178419843 | 1.99852715 |
| Map2k7 | 0.526592286 | 1.99799341 |
| Nae1 | 0.168785391 | 1.99312765 |
| Klra1 | 1.624114429 | 1.99223099 |
| Ackr1 | 0.686322356 | 1.99113229 |
| Gsk3b | 0.139950878 | 1.98458808 |
| Ddx5 | 0.108357222 | 1.98452123 |
| Rasgrp4 | -0.30796855 | 1.98204259 |
| Cd79b | 0.573624698 | 1.98181291 |
| Selenos | -0.094135389 | 1.98034559 |
| Lrrk2 | -0.279130826 | 1.97320552 |
| Ctsg | 0.772144851 | 1.97149761 |
| Csf3 | 0.879835234 | 1.97102669 |
| Crebbp | 0.155109664 | 1.96630245 |
| Ap1m1 | -0.168071183 | 1.95339155 |
| Ccl17 | 1.121472055 | 1.94998937 |
| Ifne | 0.99263584 | 1.94843066 |
| Cblb | 0.27196105 | 1.94757676 |
| Gzmk | 0.567098789 | 1.93918691 |
| Ace | -0.379916547 | 1.93625608 |
| Chuk | 0.151051318 | 1.92945839 |
| Cxcr6 | 0.524971233 | 1.92829544 |
| Hmgb1 | 0.558632621 | 1.91308922 |
| Ctla4 | 0.846563016 | 1.90031464 |
| Map3k7 | 0.358967525 | 1.89152854 |
| Ccr7 | 0.432548114 | 1.89088758 |
| Il7 | 0.871264663 | 1.89042871 |
| Nampt | -0.369920923 | 1.88630628 |
| Ifnb1 | 0.813546238 | 1.88528098 |
| Traf3 | -0.151146962 | 1.87973438 |
| Mdfic | -0.31548035 | 1.87888731 |
| Il17d | 0.503941318 | 1.87803696 |
| Ptpn4 | 0.434533448 | 1.86461496 |
| Il24 | 1.69169583 | 1.86347551 |
| Mapk1 | 0.159396537 | 1.84522333 |
| Rnasel | 0.40100559 | 1.83445649 |
| Uba52 | 0.16900681 | 1.82467436 |
| Ccr6 | 0.60857476 | 1.81838783 |
| Il1r1 | 0.386940055 | 1.81467842 |
| Nlrp3 | 0.38370467 | 1.81311349 |
| Nfat5 | 0.258162841 | 1.80605188 |

| | | |
|---|---|---|
| Tpsb2 | 0.584267083 | 1.8055031 |
| Nfkb2 | 0.173745986 | 1.801729 |
| Sp100 | -0.232311792 | 1.7995428 |
| Cd276 | 0.546086142 | 1.79490145 |
| Il1a | 0.395320689 | 1.79076597 |
| Msra | 0.446424188 | 1.78739403 |
| Il18 | -0.261548719 | 1.7810165 |
| Il18bp | -0.795412138 | 1.78061057 |
| Panx1 | 0.517773136 | 1.78033978 |
| Atf4 | 0.275874917 | 1.77597045 |
| Pik3ap1 | 0.297545008 | 1.77186613 |
| Ifnz | 0.74359002 | 1.77143327 |
| Il11ra1/2 | 0.320074032 | 1.76913711 |
| Il1r2 | 0.560648585 | 1.76793376 |
| Furin | 0.400300382 | 1.76093184 |
| Ephx2 | 0.396337866 | 1.75190081 |
| Il1f6 | 0.399076735 | 1.74866106 |
| Atf6 | 0.159191026 | 1.74764573 |
| Fcer1a | 1.137297395 | 1.736697 |
| Smad4 | 0.122052787 | 1.73630217 |
| Il1rapl1 | 0.742913548 | 1.73538174 |
| Il17rc | 0.674060916 | 1.73436311 |
| H2-Ob | 0.378374694 | 1.7268594 |
| Irak3 | 0.252322007 | 1.72432166 |
| Rps6kb1 | 0.26300508 | 1.7236856 |
| Icos | 0.704587222 | 1.72290218 |
| Ifna4 | 0.887185015 | 1.72248166 |
| Ptprc | -0.17014133 | 1.71940969 |
| Tnfrsf17 | 0.588420992 | 1.7081916 |
| Tnfsf4 | 0.54263401 | 1.70730267 |
| Il1rapl2 | 0.671901898 | 1.70649362 |
| Il6 | 0.885954072 | 1.69954378 |
| Pgk1 | -0.328147696 | 1.69884579 |
| Rela | 0.364761397 | 1.69665537 |
| Cxcl1 | 0.918662075 | 1.69573685 |
| Cmklr1 | -0.265084933 | 1.69079579 |
| Irf1 | -0.480989647 | 1.68908199 |
| Tbk1 | -0.126443577 | 1.68894513 |
| Il1f10 | 0.707879315 | 1.68688133 |
| Ifih1 | 0.219412977 | 1.68503542 |
| Plscr2 | 0.496353249 | 1.68481768 |

| | | |
|---|---|---|
| Il15 | -1.258458147 | 1.68398195 |
| Marcks | 0.48081411 | 1.66678892 |
| Cxcl2 | 0.649486041 | 1.66268925 |
| Osm | 0.466207907 | 1.66222385 |
| Ifnl2/3 | 0.683527167 | 1.65658983 |
| Sh2d1a | 0.954242509 | 1.65558036 |
| Ifnk | 0.919893266 | 1.65359314 |
| Tlr8 | -0.37727557 | 1.65179856 |
| Atg13 | 0.21792193 | 1.64881733 |
| Cxcl15 | -0.437586243 | 1.6485042 |
| Stat2 | -0.247645717 | 1.64406616 |
| Mavs | 0.337491456 | 1.64049619 |
| Il17a | 0.630001623 | 1.63887321 |
| Pik3r6 | 0.191318022 | 1.63824441 |
| Klrd1 | 0.491646886 | 1.63743364 |
| Cxcl3 | 0.901808694 | 1.63619165 |
| Txk | 0.787693512 | 1.63360316 |
| Ptger4 | 0.397962967 | 1.62856241 |
| Nrde2 | 0.57890817 | 1.62731189 |
| Nfatc1 | 0.185147579 | 1.62702228 |
| Ccl22 | 0.981946065 | 1.62697827 |
| Jaml | 0.363209645 | 1.62175068 |
| Tbp | 0.214498363 | 1.61736038 |
| Fcgr1 | -0.515965557 | 1.61275339 |
| Nfkb1 | 0.264899274 | 1.60613825 |
| Dnajc10 | 0.410362265 | 1.6056321 |
| Tlr5 | 0.490897234 | 1.58166794 |
| Ifnlr1 | 0.471471914 | 1.58113418 |
| Bcl2 | 0.567632186 | 1.57945494 |
| Xcr1 | 0.949584124 | 1.57504663 |
| Atf2 | 0.252249383 | 1.5727961 |
| Il25 | 0.445262691 | 1.57279311 |
| Kir3dl1/2 | 0.751281133 | 1.56919809 |
| Sem1 | 0.224845312 | 1.56591734 |
| Npc2 | -0.180373819 | 1.55591127 |
| Ltbr | 0.325109974 | 1.55436699 |
| Trim26 | 0.243381713 | 1.54146412 |
| Il1rl1 | 0.23478133 | 1.54075682 |
| Sigirr | 0.5478625 | 1.53882439 |
| Tnfrsf4 | 0.807849215 | 1.53707868 |
| Cd80 | 0.688823813 | 1.52333757 |

| | | |
|---|---|---|
| Tnfrsf10b | 0.656423826 | 1.52306974 |
| Lyn | -0.280655289 | 1.52245505 |
| Ifna1/5/6/12/13 | 0.750049725 | 1.52067998 |
| Irf7 | -0.615323507 | 1.51570826 |
| Ifi27 | 0.287992153 | 1.51293967 |
| Ncf2 | 0.18093307 | 1.5044876 |
| Ccl19 | 0.439364328 | 1.48853412 |
| Il18r1 | 0.238598056 | 1.48564268 |
| C3 | -0.272574233 | 1.47964519 |
| Ackr3 | 0.902257166 | 1.47775783 |
| Aif1 | -0.833285721 | 1.47540606 |
| Cd14 | 0.397518756 | 1.47031584 |
| Cx3cr1 | 0.388483899 | 1.4605226 |
| Jak2 | 0.123015853 | 1.46021683 |
| Psmb10 | -0.358130642 | 1.45953565 |
| Sell | -0.286778866 | 1.4571856 |
| Ndufs8 | 0.08360241 | 1.45100138 |
| Cx3cl1 | 0.762108422 | 1.44662332 |
| Cul1 | 0.11895102 | 1.44537894 |
| Lcn2 | 0.51048228 | 1.44271466 |
| Psmb8 | -0.330945255 | 1.43972315 |
| Xbp1 | 0.231319601 | 1.43498596 |
| Isg15 | -0.433194488 | 1.4347509 |
| Il12b | 0.478336649 | 1.4343738 |
| Ccl11 | 0.859854966 | 1.43233087 |
| Lyz2 | -0.317444766 | 1.43125725 |
| Il6st | 0.323219767 | 1.422226 |
| Ets1 | 0.599684343 | 1.41954697 |
| Marco | -0.659002124 | 1.4193002 |
| Ms4a7 | -0.536721663 | 1.41830878 |
| Kdm6b | 0.271686975 | 1.41733413 |
| P2rx7 | -0.2883666 | 1.41632347 |
| Samhd1 | -0.324638153 | 1.41394754 |
| Itgb7 | -0.510626646 | 1.39226473 |
| Ddost | -0.200603128 | 1.39127153 |
| Csf3r | -0.20432581 | 1.38718821 |
| Il17f | 0.570182272 | 1.38165307 |
| Lilra5 | 0.46658723 | 1.38065529 |
| Tnfrsf18 | 0.37640598 | 1.37793578 |
| Cxcl10 | -0.358854497 | 1.37499587 |
| Cd81 | 0.240302806 | 1.37328083 |

| | | |
|---|---|---|
| Gbp2 | -0.424588288 | 1.3617035 |
| Il22ra1 | 0.647728837 | 1.36003845 |
| Gzmn | 0.843744604 | 1.35883484 |
| Card11 | -0.128487474 | 1.35351774 |
| C3ar1 | -0.571035232 | 1.35240077 |
| Agt | 0.484238024 | 1.35203652 |
| Il13 | 0.876766423 | 1.35117735 |
| Cd40lg | 0.676379117 | 1.35059758 |
| Ppia | 0.113144609 | 1.34938449 |
| Gata3 | 0.575258528 | 1.3469947 |
| Ssr1 | 0.146565429 | 1.34312119 |
| Anpep | -0.404017353 | 1.34079597 |
| Tgfb3 | 0.620456117 | 1.34037624 |
| Plat | 0.593017993 | 1.34027728 |
| Bst2 | -0.427770143 | 1.33419782 |
| Dnaja2 | 0.155656461 | 1.33272993 |
| Ddit3 | -0.341058135 | 1.32564776 |
| Cflar | -0.146836625 | 1.32415795 |
| Os9 | -0.075896862 | 1.3234235 |
| Ccl4 | -0.278094834 | 1.31993406 |
| Tcl1 | 0.776800144 | 1.31905941 |
| Rgma | 0.452737044 | 1.31446409 |
| Hsd11b1 | -0.249734645 | 1.31348107 |
| Sele | 0.586873963 | 1.31162768 |
| Runx3 | 0.337881616 | 1.31085873 |
| Hdc | 0.459160559 | 1.30982955 |
| Ifna2 | 0.797588993 | 1.30911979 |
| Acsl4 | 0.173767518 | 1.30882799 |
| Ccr4 | 0.604666796 | 1.30880669 |
| Nlrp1a | -0.960811479 | 1.30568954 |
| Il23r | 0.535482349 | 1.30326098 |
| Il9 | 0.593407205 | 1.3011304 |
| Il2 | 0.675113315 | 1.3006782 |
| Nfatc3 | 0.150125605 | 1.29962408 |
| Gpx7 | 0.489030579 | 1.29899127 |
| Lax1 | 0.41517325 | 1.29808174 |
| Tcf7 | 1.180455213 | 1.29757405 |
| Il17rb | 0.70943437 | 1.29635317 |
| Mif | 0.28322773 | 1.2932086 |
| Il7r | -0.208494514 | 1.29212626 |
| Il31 | 0.961830135 | 1.28894807 |

| | | |
|---|---|---|
| Ccr8 | 0.73655391 | 1.28251075 |
| Psmb9 | -0.372298063 | 1.28144731 |
| Ccl7 | -0.449412889 | 1.27788033 |
| Tpsab1 | 0.647659261 | 1.27559077 |
| Tab1 | 0.30940002 | 1.27509741 |
| Cd163 | 0.532886491 | 1.27466941 |
| Lancl1 | 0.369828648 | 1.27320158 |
| Lef1 | 0.664753292 | 1.27300721 |
| Il23a | 0.347411675 | 1.26653094 |
| Mafb | -0.210587432 | 1.26416076 |
| Plau | -0.579531225 | 1.2605015 |
| Cd2 | 0.436975313 | 1.26029739 |
| Il3 | 0.692149192 | 1.25732215 |
| Eif2ak2 | -0.131004631 | 1.25153434 |
| Cxcr3 | 0.564165275 | 1.25120988 |
| Fcgr4 | -0.601220754 | 1.24677735 |
| Gzmd | 1.207974587 | 1.24321145 |
| Pdcd1lg2 | -0.11469712 | 1.24109523 |
| Ccnc | 0.090803008 | 1.23362685 |
| Lgals3 | -0.206616029 | 1.2328171 |
| Il9r | 0.677379717 | 1.22241421 |
| Ulk2 | 0.325185124 | 1.22151255 |
| Cd28 | 0.906153842 | 1.22051481 |
| Il17b | 0.563145343 | 1.21969156 |
| Plg | 0.522969316 | 1.21860331 |
| Ptpn6 | -0.234917615 | 1.21797524 |
| Il17rd | -0.660768389 | 1.21505126 |
| Akt2 | 0.155737208 | 1.21470661 |
| Tnfsf9 | 0.288736913 | 1.21255735 |
| Tln1 | -0.16177144 | 1.20200392 |
| Defb14 | 0.66022507 | 1.19675152 |
| Ltc4s | 0.268344529 | 1.1966532 |
| Lta4h | 0.177684206 | 1.19263749 |
| Cd19 | 0.462208711 | 1.19153943 |
| Il20rb | 0.338830735 | 1.18450408 |
| Pik3r5 | -0.195451242 | 1.18343193 |
| Tgfb2 | 0.359062724 | 1.17567081 |
| Mvp | -0.127567918 | 1.1700142 |
| Pdcd1 | 0.449405192 | 1.16767646 |
| Sort1 | 0.235141517 | 1.16411115 |
| Ccl26 | 0.254824962 | 1.16353158 |

| | | |
|---|---|---|
| Apbb1ip | -0.117484323 | 1.1630816 |
| Cxcl11 | -0.378593821 | 1.16133774 |
| Eomes | 0.996622694 | 1.15747041 |
| Smad3 | -0.301331484 | 1.15594654 |
| Il10ra | 0.194421457 | 1.15362311 |
| Zbp1 | -0.326898918 | 1.15094382 |
| Rnf31 | 0.192992072 | 1.14977267 |
| Casp3 | 0.141186693 | 1.14966964 |
| Dtx3l | -0.235612108 | 1.1489836 |
| Ido1 | 0.611883326 | 1.14775297 |
| Fasl | 0.326900647 | 1.14503494 |
| C5ar1 | 0.224673127 | 1.14251406 |
| Lat2 | -0.244879585 | 1.13992723 |
| Il34 | 0.498419813 | 1.1324797 |
| Pxn | 0.175065834 | 1.13044998 |
| Evl | -0.396440166 | 1.12963569 |
| Ncf4 | -0.19066215 | 1.12775775 |
| Trat1 | 1.342220425 | 1.12660649 |
| Neu1 | 0.120962171 | 1.12189649 |
| Ccr9 | 0.884478068 | 1.11803738 |
| Ripk1 | -0.246313339 | 1.11703939 |
| Syk | 0.147876976 | 1.11681188 |
| Itpr3 | 0.438271476 | 1.11253364 |
| Vwf | -0.265400521 | 1.11187397 |
| Ifitm3 | -0.202401588 | 1.1101576 |
| Adgre5 | 0.194085685 | 1.10789581 |
| Hc | -0.413048899 | 1.1066918 |
| Tnfsf13b | 0.22482449 | 1.10470498 |
| Thop1 | 0.294574142 | 1.10310453 |
| Tlr1 | -0.319102066 | 1.10020702 |
| Tcirg1 | -0.118251554 | 1.09762219 |
| Ceacam3 | 0.49834035 | 1.09722822 |
| Cd22 | 0.210353947 | 1.09363963 |
| H2-Pa | 0.663577264 | 1.0933869 |
| Cd274 | -0.230699363 | 1.09081354 |
| Plekha1 | 0.105506298 | 1.08843608 |
| Il10 | 0.497574167 | 1.08762873 |
| Lag3 | 0.392311002 | 1.08746372 |
| Pirb | -0.247588764 | 1.08731802 |
| Il22 | 1.061877891 | 1.0846646 |
| Tnfrsf25 | 0.274615521 | 1.08175367 |

| | | |
|---|---|---|
| Pik3r3 | 0.402124027 | 1.07307625 |
| Cd3e | 0.713565268 | 1.07114459 |
| Cd79a | 0.291467769 | 1.07081088 |
| Cgas | -0.222097969 | 1.07028385 |
| Mx1 | -0.368477376 | 1.06642858 |
| Alox12 | 0.228679296 | 1.06488227 |
| Maf | -0.417804346 | 1.06369633 |
| Cd70 | 0.496520413 | 1.06071743 |
| Nfatc4 | 0.397891412 | 1.05911649 |
| Cd38 | -0.308736533 | 1.05131911 |
| Plcg1 | 0.482194986 | 1.04990889 |
| Ccr3 | 0.287151862 | 1.04587364 |
| Itgb2 | -0.21830292 | 1.03307432 |
| Il21 | 0.53900764 | 1.02621319 |
| Trim33 | 0.111859392 | 1.02167363 |
| Alpl | 0.500243328 | 1.01873824 |
| H2-DMb2 | 0.253403197 | 1.01625437 |
| Neo1 | 0.322898248 | 1.01596304 |
| Cd4 | 0.70462625 | 1.01265544 |
| Mme | 0.356552885 | 1.01041778 |
| Il1rap | 0.197592689 | 0.99766731 |
| Mlkl | -0.22579979 | 0.99727761 |
| Ccl5 | -0.338566896 | 0.99464996 |
| Gzmc | 0.603106625 | 0.99357048 |
| Atm | 0.25719455 | 0.98649862 |
| Tbxas1 | -0.256761765 | 0.98589845 |
| Ltb | 0.121847046 | 0.98273866 |
| Hlx | 0.176980267 | 0.98022067 |
| H2-Q1 | 0.317976914 | 0.9800919 |
| Adar | -0.103764074 | 0.97831585 |
| Cd247 | 0.758319327 | 0.97818417 |
| Pik3cb | 0.100948107 | 0.97607206 |
| Ccl21a/b/c/d | 0.380366622 | 0.97607102 |
| Ltf | 0.479144576 | 0.97509953 |
| Gns | -0.140134586 | 0.97101076 |
| C2 | -0.332434941 | 0.96550324 |
| Vrk3 | -0.107474191 | 0.95988962 |
| Ebi3 | 0.217321815 | 0.95237274 |
| Ticam1 | 0.177288373 | 0.9488742 |
| Lamp2 | -0.131245669 | 0.94869978 |
| Oas3 | -0.132592648 | 0.93982288 |

| | | |
|---|---|---|
| Eif2ak3 | 0.448786052 | 0.93949146 |
| Ern1 | -0.186338518 | 0.93857055 |
| Vcam1 | 0.338628344 | 0.9340506 |
| Il11 | 0.57368804 | 0.93026151 |
| Cd3d | 0.537783114 | 0.92825453 |
| Klrk1 | -0.325505937 | 0.928096 |
| Il2rb | 0.236341233 | 0.92664552 |
| Gbp3 | -0.240765499 | 0.91642177 |
| Cd8a | 0.769742385 | 0.91440607 |
| Il1rl2 | 0.163408527 | 0.91085472 |
| H2-T23 | -0.304734378 | 0.90965397 |
| Acvr1 | 0.273154156 | 0.90699228 |
| Ms4a1 | 0.34038156 | 0.90576621 |
| Itgae | 0.541877148 | 0.90484717 |
| Cbfb | -0.122605579 | 0.90374523 |
| Ccl1 | 0.341531442 | 0.89669951 |
| Cd86 | 0.1380872 | 0.88578683 |
| Cd59a | 0.515145592 | 0.88512555 |
| Tab2 | 0.099290513 | 0.88269115 |
| Prkcsh | 0.07813726 | 0.87958564 |
| Batf | 0.225080873 | 0.87259018 |
| Cd96 | 0.504129232 | 0.87087537 |
| Alpk1 | -0.168763878 | 0.86354662 |
| Ccr1 | 0.180673966 | 0.86289663 |
| Ifi44 | 0.217750772 | 0.86217065 |
| Bdkrb2 | 0.276975276 | 0.8616013 |
| Gzmm | 0.2810113 | 0.85774645 |
| Glb1 | 0.347538135 | 0.85460359 |
| Il20ra | 0.298354697 | 0.85044403 |
| Cxcr4 | 0.108067863 | 0.84669715 |
| Trim5 | 0.277576807 | 0.84591547 |
| Zap70 | 0.360679089 | 0.84576385 |
| Csf1r | -0.10867108 | 0.83475141 |
| Irf3 | -0.121237623 | 0.83430772 |
| Ccl12 | -0.569654402 | 0.82923689 |
| Spib | 0.303153471 | 0.82807495 |
| Prkcd | 0.074448924 | 0.82605982 |
| Sp1 | 0.070849626 | 0.82362074 |
| Gzma | -0.456751013 | 0.82292015 |
| Il27 | -0.433595837 | 0.81623704 |
| Pik3c3 | 0.1772638 | 0.81180576 |

| | | |
|---|---|---|
| Plaur | 0.353075673 | 0.81145567 |
| Cbl | 0.085713118 | 0.8019294 |
| Ddah2 | 0.187091033 | 0.80140185 |
| Tlr9 | 0.138847294 | 0.79672514 |
| Lamp3 | 0.109142204 | 0.79199587 |
| Ripk3 | 0.129094953 | 0.78030795 |
| Gadd45b | -0.105408599 | 0.77644823 |
| Sting1 | -0.231561573 | 0.77583164 |
| Fcgrt | 0.207650297 | 0.77361981 |
| Ifng | -0.513352001 | 0.77015672 |
| Ifit2 | -0.222857607 | 0.76739303 |
| Ccl25 | 1.239653912 | 0.76362124 |
| Il31ra | 0.233731487 | 0.75800633 |
| Cebpb | 0.093498535 | 0.75767972 |
| Rnf114 | -0.11945108 | 0.75452905 |
| Itk | 0.941462024 | 0.74746502 |
| Ifngr2 | -0.065506487 | 0.74588218 |
| Tlr4 | -0.114781622 | 0.74254988 |
| Casp4 | -0.139161938 | 0.73968338 |
| Map2k2 | 0.107567623 | 0.73927169 |
| Akt3 | 0.120491362 | 0.73897093 |
| Crcp | -0.13517159 | 0.729061 |
| Gucy1a1 | 0.233788717 | 0.72560391 |
| Prkcq | 0.391902158 | 0.72067676 |
| Casp1 | -0.139671256 | 0.71727221 |
| Tyk2 | 0.148002755 | 0.71178142 |
| Oaz1 | -0.084727438 | 0.70137982 |
| Vsir | 0.114210497 | 0.700608 |
| Cd6 | 0.601836277 | 0.69845978 |
| Diablo | -0.095206035 | 0.69473486 |
| Cd40 | -0.264256848 | 0.69413363 |
| Cxcl12 | -0.236084863 | 0.69201006 |
| Ifngr1 | 0.104844302 | 0.69067752 |
| Bnip3 | 0.197242058 | 0.68386484 |
| Bcl3 | 0.111006459 | 0.68191388 |
| Lck | 0.437928838 | 0.67922197 |
| Smad5 | 0.093852119 | 0.67852841 |
| Il15ra | -0.146112551 | 0.6697133 |
| Csf2rb | 0.105401295 | 0.66960302 |
| Cd8b1 | 0.571721525 | 0.66070443 |
| Dysf | 0.382144997 | 0.65890777 |

| | | |
|---|---|---|
| Tirap | 0.152694774 | 0.658243 |
| Nmt1 | -0.111047446 | 0.65500207 |
| Stt3b | 0.128820627 | 0.65266486 |
| Pnoc | 0.493134223 | 0.65253907 |
| Rab5c | -0.137372035 | 0.6480066 |
| App | 0.088833534 | 0.64593936 |
| Itgam | -0.206000583 | 0.64529833 |
| Il5 | 0.478753584 | 0.6442194 |
| Il21r | 0.144017465 | 0.63781838 |
| Thbs1 | 0.28630228 | 0.63542224 |
| Cxcl9 | -0.533963912 | 0.62975741 |
| Psen1 | 0.07467663 | 0.62944337 |
| Ifit3 | 0.111302897 | 0.62928378 |
| Rac2 | 0.123144137 | 0.62754424 |
| Irf9 | 0.080514469 | 0.62736199 |
| Ccl2 | -0.189783372 | 0.62558081 |
| Nlrc4 | 0.161402052 | 0.62471636 |
| Hprt | 0.058423778 | 0.62010395 |
| Il13ra1 | -0.09895268 | 0.61568832 |
| Trim21 | -0.155122173 | 0.60081288 |
| Parp1 | -0.190630918 | 0.59100954 |
| Il17c | 0.517295452 | 0.58506046 |
| Csf2ra | 0.117302662 | 0.58420624 |
| Slc11a1 | -0.096177212 | 0.57543559 |
| Mcl1 | 0.096541556 | 0.57335528 |
| Ikbkb | 0.095343236 | 0.56885816 |
| Nod2 | 0.113294494 | 0.56729518 |
| Fcrlb | -0.160092388 | 0.56712712 |
| Txn1 | -0.132035524 | 0.56497881 |
| Rab7 | 0.119416336 | 0.56466441 |
| Map3k5 | -0.039806324 | 0.55233906 |
| Rps6ka3 | -0.062599741 | 0.5505233 |
| Mgam | 0.177999713 | 0.55029605 |
| Cd36 | -0.083629258 | 0.55017662 |
| Raf1 | 0.082417536 | 0.54894014 |
| Stat6 | 0.087060608 | 0.54838453 |
| Atg12 | 0.071738809 | 0.54204844 |
| Il4 | 0.183736023 | 0.53770101 |
| Nod1 | -0.155927288 | 0.53314413 |
| Ifitm1 | -0.192695274 | 0.5324727 |
| Stat3 | 0.055776089 | 0.52875076 |

| | | |
|---|---|---|
| Ncf1 | 0.078379148 | 0.52377867 |
| Mtor | 0.100665518 | 0.52291756 |
| Stat5b | -0.079240606 | 0.51723001 |
| Slc2a3 | 0.125682342 | 0.51604007 |
| Pik3ca | 0.070453642 | 0.50872442 |
| Mefv | -0.070069404 | 0.50590873 |
| Bpi | 0.191045397 | 0.50535342 |
| Tap2 | -0.130422103 | 0.50324076 |
| Irf4 | 0.156532349 | 0.50111154 |
| Peli1 | -0.071403003 | 0.49952377 |
| Tgfb1 | 0.092912918 | 0.49576304 |
| Lat | 0.479454671 | 0.49336081 |
| Pstpip1 | 0.071972562 | 0.49056187 |
| Il27ra | -0.273602886 | 0.48704757 |
| Il33 | -0.157839736 | 0.48540783 |
| Ncr1 | 0.182799213 | 0.48456888 |
| Atg4a | 0.081979998 | 0.48033933 |
| Itgax | 0.092146274 | 0.47262701 |
| Pycard | -0.107589363 | 0.4682516 |
| Pik3cd | 0.07489626 | 0.4657418 |
| Rhog | -0.071377974 | 0.4647479 |
| Alas1 | 0.051777013 | 0.4590673 |
| Lilra6 | -0.091613895 | 0.45469861 |
| Was | -0.053133208 | 0.45279064 |
| Ackr2 | 0.195024859 | 0.44935319 |
| Gzme | 0.215583003 | 0.44680242 |
| Hsp90ab1 | -0.099169214 | 0.44627994 |
| Il22ra2 | -0.183985612 | 0.44213021 |
| Tyrobp | -0.062819049 | 0.4403956 |
| Mapk14 | 0.056552525 | 0.43321188 |
| Rack1 | 0.036225933 | 0.43297389 |
| Ptgs2 | 0.233601537 | 0.43173043 |
| Klrb1 | 0.297137586 | 0.43006067 |
| Cfd | 0.207121194 | 0.41868811 |
| Ms4a4a | -0.166763536 | 0.41015448 |
| Cpa3 | 0.126047024 | 0.40770292 |
| Nos2 | -0.150841044 | 0.40753404 |
| Rbck1 | 0.080032031 | 0.40453758 |
| Derl1 | -0.037522826 | 0.4025521 |
| Cap1 | -0.061541876 | 0.40098505 |
| Tlr7 | 0.071661348 | 0.39702228 |

| | | |
|---|---|---|
| Fpr1 | -0.116594307 | 0.39181398 |
| Il2ra | -0.111204868 | 0.38639685 |
| Entpd1 | -0.103930987 | 0.38218826 |
| Acsl1 | 0.073360301 | 0.38021304 |
| Cxcl13 | 0.148620211 | 0.37907574 |
| Rasgrp1 | 0.131456355 | 0.37565527 |
| Tbx21 | 0.134444329 | 0.37437594 |
| Vegfa | 0.046302627 | 0.36964501 |
| Hpgd | -0.117730006 | 0.36143934 |
| Ccl8 | -0.339753615 | 0.36029727 |
| Ccrl2 | 0.048486361 | 0.35559038 |
| Pfkfb3 | 0.070552267 | 0.35555445 |
| Fcgr2b | 0.089558207 | 0.34957072 |
| Ccl9 | 0.083243525 | 0.33248834 |
| Plek | 0.061991734 | 0.33139235 |
| Il1b | 0.130141249 | 0.33040042 |
| Abcf1 | 0.046714402 | 0.32643717 |
| Cxcr2 | 0.103700398 | 0.32086399 |
| Nkg7 | 0.108552945 | 0.32084473 |
| Cdk4 | -0.038749664 | 0.31152319 |
| Ctsl | 0.073997604 | 0.30471243 |
| Bdkrb1 | 0.103810049 | 0.30171668 |
| Sod2 | -0.06712035 | 0.30114473 |
| Ngly1 | 0.049375407 | 0.29785893 |
| Trim25 | 0.0264586 | 0.29435049 |
| Lcp2 | -0.049592883 | 0.2908211 |
| Hck | -0.057752896 | 0.2900884 |
| Fgr | -0.067557461 | 0.28888252 |
| Ube2l6 | -0.07496026 | 0.26768236 |
| Ldhb | 0.065684026 | 0.26754291 |
| Il12rb2 | 0.114201671 | 0.26599138 |
| Ifi35 | -0.068740458 | 0.26593961 |
| Ccl28 | 0.106324144 | 0.26419299 |
| Cd69 | -0.067572494 | 0.25712029 |
| Egln1 | -0.027924262 | 0.25122031 |
| Arrb2 | 0.068479583 | 0.25055016 |
| Tap1 | -0.045711734 | 0.24770481 |
| Tmem140 | 0.11185769 | 0.24546663 |
| H2-K1 | -0.078697934 | 0.24319431 |
| Timp2 | 0.033854205 | 0.24241453 |
| Il3ra | -0.052544156 | 0.24203063 |

| | | |
|---|---|---|
| Jun | 0.043692853 | 0.24035482 |
| Atg10 | 0.091166436 | 0.2399315 |
| Fyn | -0.08149041 | 0.23617366 |
| Tlr2 | 0.041737213 | 0.2342791 |
| H2-M3 | -0.068995781 | 0.2243025 |
| Xcl1 | -0.175632787 | 0.2196005 |
| Il13ra2 | 0.107410964 | 0.21858275 |
| Itln1 | 0.116126406 | 0.21542915 |
| Hsp90aa1 | -0.040090246 | 0.20991469 |
| Lrg1 | 0.045366152 | 0.20199499 |
| Il12rb1 | 0.081154041 | 0.20118159 |
| Il20 | -0.245683897 | 0.19847137 |
| Il17re | 0.076624194 | 0.19768765 |
| Tram1 | 0.028176959 | 0.19616165 |
| Ctsw | 0.052026297 | 0.1949993 |
| Gk | 0.050999211 | 0.19249498 |
| Ddx58 | -0.03852529 | 0.19023258 |
| H2-DMb1 | -0.058798884 | 0.18769246 |
| Dhx58 | -0.044437561 | 0.17755558 |
| Tlr6 | 0.026357389 | 0.17701738 |
| Blk | 0.060857257 | 0.17520994 |
| Prkca | 0.066583619 | 0.1751087 |
| Alox5ap | 0.0418051 | 0.17356521 |
| Lcp1 | -0.040080578 | 0.16811132 |
| Alox15 | -0.051958199 | 0.16667292 |
| Pecam1 | 0.029676739 | 0.16337409 |
| Socs1 | 0.058917701 | 0.16183361 |
| Ripk2 | 0.021952381 | 0.15986334 |
| Ccl6 | 0.043351299 | 0.15747641 |
| Mapk9 | -0.028869565 | 0.1572284 |
| Rnf135 | 0.049120737 | 0.15503707 |
| Ifnar2 | -0.026129026 | 0.15501755 |
| Calm1 | 0.03266372 | 0.14479481 |
| Myd88 | 0.030055409 | 0.13620617 |
| Cxcl16 | -0.039980853 | 0.1328329 |
| Kpnb1 | -0.017296168 | 0.13239095 |
| Acsl3 | 0.028619736 | 0.1319734 |
| Bcl2l1 | -0.02066578 | 0.13130338 |
| Ifnar1 | -0.015707727 | 0.13016929 |
| Rab31 | 0.042257106 | 0.12965519 |
| Cxcr1 | 0.038917843 | 0.12867884 |

| | | |
|---|---|---|
| Fpr2 | -0.04594497 | 0.12692828 |
| Rsad2 | -0.042255441 | 0.12599227 |
| Nras | 0.01635667 | 0.12561438 |
| Cd3g | 0.065977919 | 0.12329025 |
| Fbxo6 | -0.016036233 | 0.11484286 |
| Ap1g1 | 0.013331024 | 0.11475831 |
| H2-Ab1 | 0.052526713 | 0.11151687 |
| Prf1 | -0.049716592 | 0.10730685 |
| Jak1 | 0.008121155 | 0.10595927 |
| Spi1 | 0.030628104 | 0.10469002 |
| Cxcl17 | 0.03180964 | 0.1038639 |
| Xaf1 | -0.030537423 | 0.10309528 |
| Ptk2b | -0.017541457 | 0.10126173 |
| Tifa | 0.023489097 | 0.10026811 |
| Atg3 | -0.010394534 | 0.09929754 |
| Tnfsf18 | -0.065746914 | 0.09853082 |
| Gucy1b1 | -0.098179262 | 0.0980959 |
| Mknk1 | -0.034747392 | 0.09682235 |
| Ccr10 | -0.054502713 | 0.09678878 |
| Sugt1 | -0.010803213 | 0.09230409 |
| Traf2 | -0.016337703 | 0.09211279 |
| H2-Eb1 | 0.042113621 | 0.08096 |
| Ccl3 | 0.028612167 | 0.08034252 |
| Casp8 | 0.011801523 | 0.07990223 |
| Limk2 | 0.006375259 | 0.07483444 |
| Nfatc2 | -0.036567851 | 0.06903352 |
| Acox1 | -0.014710905 | 0.06147415 |
| Foxp3 | -0.031381421 | 0.05291796 |
| H2-D1 | 0.018637151 | 0.05016397 |
| Oasl1 | -0.018365505 | 0.04860263 |
| Tnfsf10 | -0.019927548 | 0.04770238 |
| Apol6 | -0.024331486 | 0.04306322 |
| Aim2 | 0.00782371 | 0.03989907 |
| Pak1 | 0.009645382 | 0.03636976 |
| Rps6ka1 | -0.007217537 | 0.03334089 |
| Ap1s2 | -0.005840391 | 0.03238679 |
| Litaf | -0.00509747 | 0.0286155 |
| Sdha | 0.007034474 | 0.01853867 |
| Tnfrsf1a | -0.002540778 | 0.01835603 |
| Parp9 | 0.004147176 | 0.0152149 |
| Jak3 | 0.003750335 | 0.01041914 |

| | | |
|---|---|---|
| Ikbke | 0.002572276 | 0.00784507 |
| Cd244a | -0.001479411 | 0.00782948 |
| Ifit1 | -0.001930659 | 0.00583671 |
| Creb1 | -0.00103292 | 0.00487958 |
| H2-DMa | -0.001352163 | 0.00270448 |
| Il1rn | 0.000564228 | 0.00141523 |
| Mrc1 | 0.000575561 | 0.00130547 |
